# Modulation of retroviral capsid assembly halts ARC-mediated TDP-43 intercellular spreading

**DOI:** 10.64898/2026.07.13.738236

**Authors:** Andrés Jiménez-Zúñiga, Alex M. Ascensión, Ángela Sánchez-Molleda, Irene Jiménez-Salvador, Martín Fuentetaja Leza, Markel G Ibarluzea, Rafael Ramis, Laura Rodríguez-Gómez, Haizea Hernández-Eguiazu, Maddi Alcain-Dominguez, José Luis Zúñiga-Elizari, Saioa Moragón, Anabel Saez-Mas, Lorea Blázquez, Vanesa Lafarga, Oscar Fernandez-Capetillo, Aritz Leonardo, Aitor Bergara, Adolfo López de Munain, Francisco Javier Gil-Bea, Gorka Gerenu Lopetegi

**Author notes:** Authors contributed equally to this work (Andrés Jiménez-Zúñiga, Alex M. Ascensión, Ángela Sánchez-Molleda). Corresponding Authors who equally contributed to this work (Adolfo López de Munain, Francisco Javier Gil-Bea, Gorka Gerenu Lopetegi).

## Abstract

Intercellular propagation of pathological TDP-43 drives the progression of ALS and FTD, yet the mechanisms enabling transmission remain elusive. Here, we demonstrate that Activity-regulated cytoskeleton associated protein (ARC/Arg3.1) is pathologically subverted to act as a retroviral-like capsid vehicle for TDP-43 spreading. In a Drosophila model, we reveal that massive Arc1 upregulation drives glia-to-neuron TDP-43 seeding, while its genetic ablation halts pathological transfer, improves motor function, and extends survival. In parallel human cellular models, stress-induced ARC colocalizes with TDP-43 to orchestrate its intercellular transmission. Guided by ARC’s capsid architecture, we reproposed Lenacapavir, an FDA-approved HIV-1 capsid modulator, as a stable binder of ARC/Arg3.1 capsid interfaces. Lenacapavir treatment effectively blocks TDP-43 propagation in vitro and rescues disease phenotypes in vivo. Our findings establish ARC as a conserved vehicle for pathological protein transmission and deliver an immediately translatable pharmacological strategy to arrest disease progression in ALS and FTD.

## 2. INTRODUCTION

Amyotrophic lateral sclerosis (ALS) and frontotemporal dementia (FTD) lie on a clinicopathological continuum characterized by a relentless, spatially progressive deterioration of motor and cognitive functions^1^. This clinical progression closely mirrors the anatomical dissemination of misfolded protein aggregates throughout the central nervous system (CNS). The defining histopathological hallmark of these neurodegenerative disorders is the intracellular accumulation of toxic protein deposits, most prominently TDP-43, but also FUS and SOD1^2,3^. Mounting evidence indicates that the progressive nature of ALS and FTD is driven by a prion-like mechanism, wherein pathological protein aggregates act as seeds that corrupt native proteins in healthy neighboring cells, thereby propagating the proteinopathy in a cell-to-cell manner^4–8^.

Despite the critical role of intercellular propagation in driving disease progression, the precise molecular vehicles orchestrating the transfer of these toxic seeds remain a major unresolved enigma. Several putative routes have been proposed based on human pathology and *in vitro* studies. Anatomical staging of post-mortem human brains reveals a sequential distribution of TDP-43 pathology (progressing from pre-inclusions to mature inclusions) along functionally connected networks, such as the monosynaptic hippocampal circuitry^9,10^. This strongly suggests a transsynaptic spread along axonal pathways, a concept further supported by the localization of pathological TDP-43 within dendritic postsynaptic densities^11,12^. Parallel studies have implicated alternative transcellular pathways, including direct cytoplasmic exchange via actin-rich tunneling nanotubes, or extracellular transmission via exosomes carrying pathogenic TDP-43 fragments^7,8,13^. However, the exact physiological relevance of these pathways remains heavily debated, particularly whether exosomes primarily serve a clearance function or act as active pathogenic vectors^14^. Ultimately, the specific identity of the dominant transport vehicles, the pathways governing their biogenesis, and the mechanisms by which they selectively package neurotoxic cargo for transmission remain largely unknown. This fundamental knowledge gap severely hinders the development of targeted therapeutic interventions capable of halting the anatomical spread of the disease.

A significant impediment to deciphering these non-cell-autonomous pathways is the scarcity of *in vivo* models specifically designed to recapitulate the intercellular transmission of proteinopathies. Most existing animal models of ALS and FTD rely on the ubiquitous or broad expression of disease-associated proteins across the CNS^15^. By forcing simultaneous pathology in virtually all cells, these models obscure the spatiotemporal segregation necessary to trace the definitive journey of a pathological seed from a specific donor population to a recipient cell^16^. Consequently, unravelling the mechanics of propagation demands *in vivo* paradigms that restrict initial protein expression to a discrete cell population. Recent advances underscore the power of this approach; for example, murine models utilizing focal viral delivery have successfully traced the anterograde and retrograde transmission of TDP-43 along specific corticospinal circuits^17^. Similarly, genetically restricted *Drosophila* models have been pivotal in demonstrating how the localized expression of TDP-43 in glial cells triggers neuronal cell death^18,19^. Implementing such paradigms by restricting initial expression specifically to glial cells, which are deeply implicated in ALS/FTD pathophysiology^20^, allows for the unequivocal tracking of pathogenic spread to neighboring neurons.

To overcome this methodological hurdle, we engineered a novel *Drosophila melanogaster* model featuring the conditional expression of human TDP-43 strictly restricted within glial cells, enabling the precise, time-dependent tracking of native transcellular seed propagation. Unbiased single-cell transcriptomic profiling of this paradigm unmasked a profound cellular transition, wherein homeostatic glia transform into a disease-associated state defined by a massive, unexpected upregulation of *Activity-Regulated Cytoskeleton Associated Protein* (*Arc1)*. *Arc1*, and its mammalian ortholog *Activity-Regulated Cytoskeleton Associated Protein (ARC)*, encodes a conserved protein derived from an endogenized retrotransposon that uniquely retains the ancestral capacity to assemble retroviral-like capsids. Expanding upon the framework recently established for tau transmission^21^, we demonstrate that under ALS/FTD proteotoxic stress, this domesticated machinery is broadly co-opted, reverting to its viral-like nature to encapsidate and propagate toxic TDP-43 seeds. This establishes the ARC capsid architecture as a convergent, cross-disease vector for neurodegenerative spreading. Crucially, we exploit this retroviral heritage as an unexpected structural vulnerability. By targeting the three-dimensional capsid assembly hub with Lenacapavir, an FDA-approved Human Immunodeficiency Virus 1 (HIV-1) capsid assembly modulator, we demonstrate that transcellular TDP-43 seeding was halted, successfully preventing downstream neuronal nuclear loss-of-function and dissemination of TDP-43 *in vivo* and *in vitro*, together with reversing motor deterioration and shortened lifespan *in vivo*. Collectively, our findings elevate ARC capsids into a universal law of neurodegenerative progression while delivering an immediate, clinically translatable strategy to arrest the anatomical spread of ALS and FTD.

## 3. RESULTS

### 1. Glia-restricted hTDP-43 expression induces transcellular spreading and functionally toxic seeding in recipient neurons

To overcome the limitations of previous expression models and trace the directional intercellular propagation of TDP-43 *in vivo*, we generated a *Drosophila melanogaster* model restricting human wild-type TDP-43 (hTDP-43) expression exclusively to glial cells using the REPO-Gal4 driver and its control counterpart (Glia-hTDP43 and Control strains; **Figure 1a**). Phenotypically, the Glia-hTDP43 strain exhibited a significant reduction in motor function at 7 days post-eclosion, along with a markedly reduced lifespan, reflecting the pathogenic impact of glia-restricted overexpression of hTDP-43 (**Figure 1b**). To characterize the temporal progression of the pathology within the glial population, we performed immunofluorescence assays on the Glia-hTDP43 strain fly brains (optic lobe and central brain) at day 1 (presymptomatic) and day 14 (late-symptomatic) post-eclosion. First, we observed early and sustained pathological changes initiated strictly within glial cells. Specifically, hTDP-43 progressively mislocalized, from the cell soma to form granular structures within glial processes by day 14 post-eclosion (**Figure 1c**). Crucially, a significant proportion of hTDP-43 became hyperphosphorylated (p-hTDP-43), the defining hallmark of pathological aggregation and progression of the pathology. This accumulation was connected with progressive apoptotic signalling, as evidenced by the concomitant rice of p-hTDP-43 and cleaved caspase-3 at 14 days of age (**Figure 1d**). Furthermore, despite compensatory activation of the ubiquitin-proteasome system and autophagy (**Figure S1a and S1b**), the glia exhibited clear signs of lysosomal dysfunction, highlighted by the accumulation of lamellar bodies (**Figure S1c**), indicating an eventual failure of the cellular clearance machinery.

**Figure 1.**
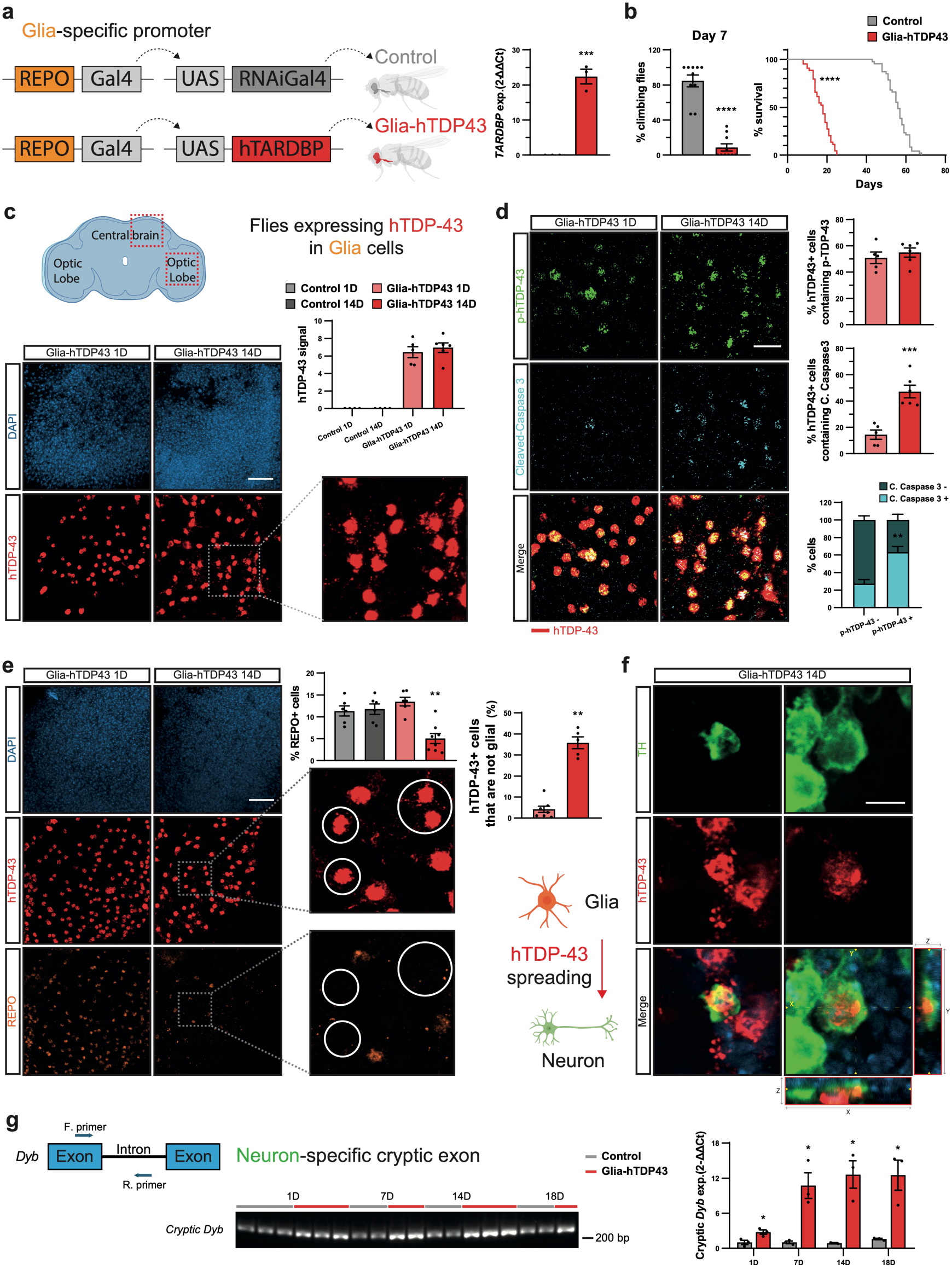
Glial expression of human TDP-43 in *Drosophila* induces ALS/FTD-like pathological features and spreading to neurons. **(a)** Schematic representation of the glia-specific driver line (REPO-Gal4) crossing strategy used to generate Control and Glia-hTDP43 models, and qPCR validation of *TARDBP* expression in whole brains. **(b)** Motor performance quantified via climbing assay at day 7 (left) and survival curves (right) demonstrating accelerated mortality in Glia-hTDP43 flies. **(c)** Schematic layout of the fly brain (central brain and optic lobes) and representative confocal projections of Glia-hTDP43 brains at 1 and 14 days post-eclosion stained with DAPI (blue) and anti–hTDP-43 (red), alongside quantification of mean hTDP-43 fluorescence signal intensity (n = 5). Scale bar = 20 μm. **(d)** Confocal images of Glia-hTDP43 brains stained with anti–hTDP-43 (red), anti–p-hTDP-43 (green), and anti–cleaved caspase-3 (cyan) at 1 and 14 days. Histograms show the quantification of (i) p-hTDP-43⁺ cells among total hTDP-43⁺ cells, (ii) cleaved caspase-3⁺ cells among total hTDP-43⁺ cells, and (iii) the proportion of cleaved caspase-3⁺ cells within p-hTDP-43⁺ versus p-hTDP-43⁻ populations at 14 days (n = 5). Scale bar = 10 μm. **(e)** Spatial tracking of non-cell-autonomous hTDP-43 spreading. Confocal images of Glia-hTDP43 brains stained with DAPI (blue), anti-REPO (orange), and anti–hTDP-43 (red) at 1 and 14 days. Graphs show the quantification of (i) the percentage of total REPO⁺ glial cells and (ii) the percentage of hTDP-43⁺ cells at 1 and 14 days (n = 6). Detail at 14 days highlights hTDP-43⁺/REPO⁻ cells. hTDP-43⁺ cells that are non-glial (REPO⁻). High-magnification details highlight intact glial restriction at day 1 and spreading into surrounding REPO⁻ cells (white circles) at day 14. Scale bar = 20 μm. **(f)** High-magnification confocal images of a 14-day-old Glia-hTDP43 brain stained with DAPI (blue), anti-TH (dopaminergic neurons, green), and anti–hTDP-43 (red), showing the internalization of glia-derived hTDP-43 withinTH⁺ neuronal populations. Scale bar = 5 μm. **(g)** Schematic layout of the *Dyb* gene locus indicating primers flanking the neuron-specific cryptic exon (left). Semi-quantitative PCR gel electrophoresis (center; 214 bp) and qPCR quantification (right) of *Dyb* cryptic exon expression in Control and Glia-hTDP43 brains at 1, 7, 14, and 18 days (n = 3). All representative images were acquired under identical optical settings and magnification. Data are shown as mean ± SEM (n = 3–6 biological replicates per group, as specified in individual panels). \**p* < 0.05, **p < 0.01, \*\*\**p* < 0.001, ****p < 0.0001 (Student’s *t*-test or Mantel-Cox log-rank test for the longevity assays).

Having established robust hTDP-43-induced pathological alterations and TDP-43 pathology within the glial compartment in this fly model, we next investigated whether these cells could act as donors for non-cell-autonomous spreading of hTDP-43. By day 14, coincidental with a marked reduction in REPO-positive glial cells, we detected a striking emergence of hTDP-43 immunoreactivity in a distinct population of non-glial cells (REPO-negative and hTDP-43-positive cells) (**Figure 1e**), suggesting that TDP-43 may spread beyond the glial cells, being the donor cells in this hTDP-43 spreading system. To confirm neurons as recipient cells, we immunolabeled dopaminergic neurons for tyrosine hydroxylase (TH) and detected hTDP-43 within their cytoplasm at day 14, a phenomenon entirely absent in younger flies (**Figure 1f**). This finding provided direct physical evidence of glia-to-neuron transcellular propagation of hTDP-43. Beyond, we sought to determine whether this intercellular transfer consisted of inert protein debris or structurally competent seeds capable of corrupting the recipient neuron’s homeostasis. Pathological hTDP-43 aggregates are known to sequester the endogenous TDP-43^22^ or its *Drosophila* ortholog Tbph^23^, leading to a catastrophic loss of nuclear function^6^. In our fly model, tracking with the spread of hTDP-43 into neurons, we observed a strong and progressive enrichment of a cryptic exon within the *Dyb* transcript, a strictly validated molecular marker for Tbph loss-of-function^24^ in *Drosophila* neuronal cells (**Figure 1g**). This finding demonstrates that hTDP-43 transferred from donor glia is capable of seeding in recipient neurons, sequestering its ortholog Tbph to drive a loss-of-function phenotype, thereby emulating the dual mechanisms of TDP-43 pathology: nuclear loss-of-function and cytoplasmic gain-of-function.

Finally, given that neurons seem to be affected by TDP-43 pathology, we sought to determine whether this impairment contributes to the motor dysfunction of the fly model. The Glia-hTDP43 model exhibited a marked structural deterioration of the presynaptic active zones (nc82-positive), resulting in a significant reduction in both synaptic bouton density and size along motor axons at day 7 of age (**Figure S1d**).

Taken together, these data establish a directional, spatiotemporal cascade whereby glial TDP-43 pathology actively spreads to adjacent neurons, seeding functional toxicity that ultimately dismantles the neuromuscular circuitry.

### 2. Progressive emergence of a disease-associated glial state culminates in massive Arc1 upregulation in response to hTDP-43

Based on the functional toxicity of glia-to-neuron hTDP-43 spreading of the fly model, we performed single-cell RNA sequencing (scRNA-seq) to identify the global transcriptional landscape underlying this non-cell-autonomous cascade. To this end, brains from both Control and Glia-hTDP43 flies were profiled across a temporal continuum: day 1 (pre-symptomatic stage), day 7 (early symptomatic stage), and day 14 (late symptomatic stage) (**Figure 2a**). To ascertain whether our findings represent a universal mechanism of ALS/FTD propagation rather than a TDP-43-specific artifact, we cross-validated our analysis using a rapidly progressing model that expresses hFUS in glia cells (Glia-hFUS, **Figure S2a-d**), with samples obtained at 1 day post-eclosion.

**Figure 2.**
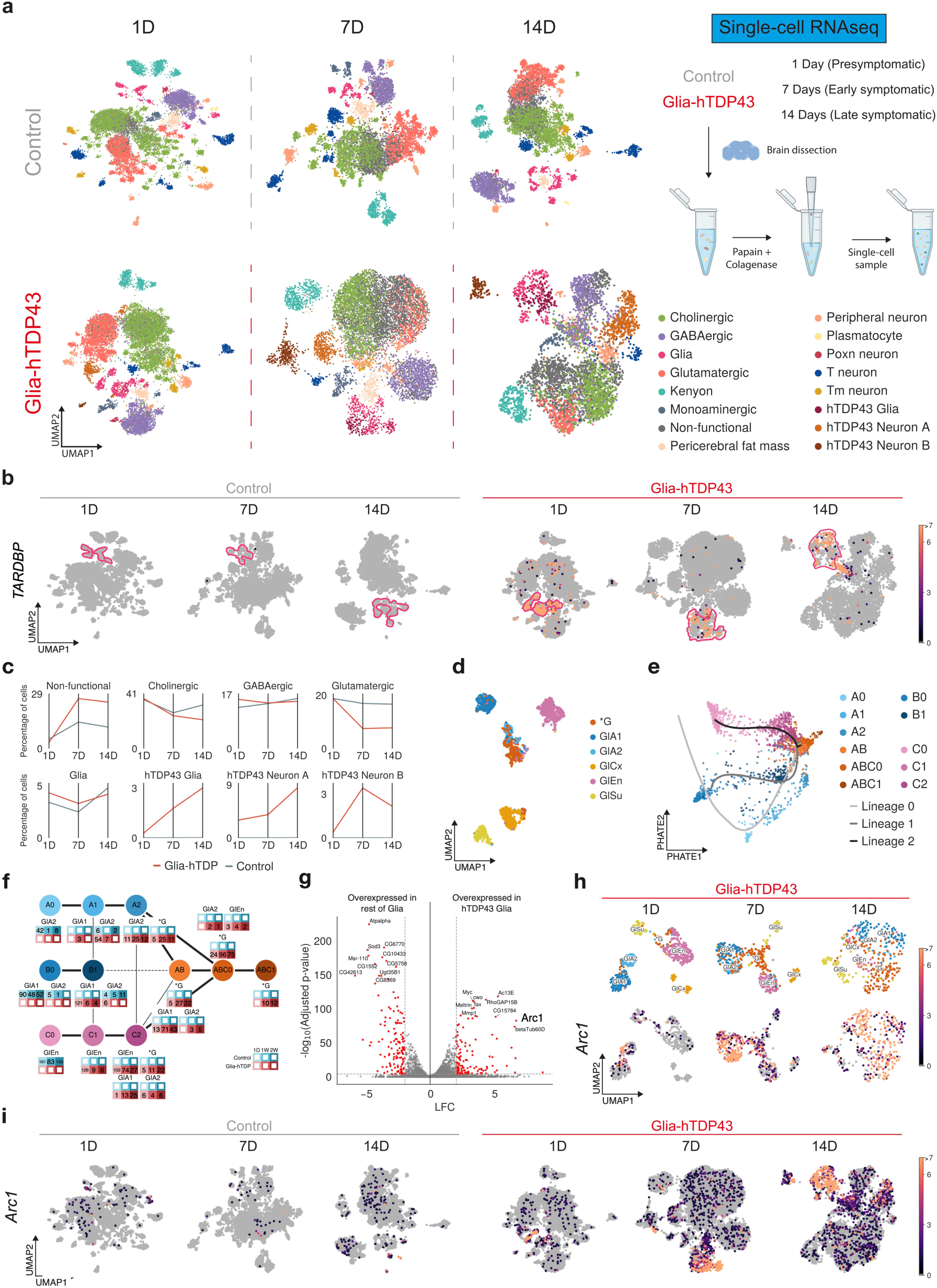
scRNA-seq reveals selective neuro-glial vulnerability and the transcriptional emergence of a pathology-associated glial subpopulation with strong *Arc1* upregulation in the Glia-hTDP43 model. **(a)** Schematic workflow of the scRNA-seq pipeline tracking *Drosophila* brains across presymptomatic (1 day), early symptomatic (7 days), and late symptomatic (14 days) stages (top right). Integrated UMAP plots (left) display all captured single-cell transcriptomes for Control and Glia-hTDP43 conditions across timepoints, color-coded by major neural and non-neural cell type identities. **(b)** Feature expression UMAP plots tracking the chronological distribution of the human *TARDBP* transgene transcript in the Glia-hTDP43 model. Note that while expression is strictly restricted to the designated glial clusters at early stages, a distinct non-glial (neuronal) subpopulation prominently contains the transcript by day 14. Glial clusters are contoured to facilitate visualization across timepoints. **(c)** Longitudinal quantification of cell population dynamics. Line graphs plot the shifting relative proportions (Δ percentage of cell) between Glia-hTDP43 and Control datasets over time, illustrating the progressive decay of specific cholinergic and glutamatergic populations alongside the synchronized emergence of a novel disease-associated glial state (*hTDP43 glia*). **(d)** UMAP plot of the baseline glial cell compartment, resolving major canonical identities: *Astrocyte-like glia* (GlA1, GlA2), *Cortex glia* (GlCx), *Ensheathing glia* (GlEn), *Surface glia* (GlSu), and transitional reactive glial subpopulation, termed as *hTDP43 Glia* (G*). **(e)** PHATE plot illustrating the pseudo-temporal evolutionary continuum of the glial population. Slingshot trajectory inference overlays outline three convergent developmental trajectories (Lineages 0, 1 and 2) migrating from peripheral homeostatic clusters (A0, B0, C0) towards a centralized, highly reactive transition cluster (ABC1). **(f)** Partition-based graph abstraction (PAGA) network showing increased connectivity (thicker edges in the graph) among the PHATE-derived clusters. Charts below each node quantify the composition of each cluster based on specific glial subtypes, chronological timepoints, and experimental conditions. **(g)** Volcano plot highlighting global differentially expressed genes (DEGs) within the glial lineage comparing Control versus Glia-hTDP43. The retroviral-like transcript *Arc1* stands out within the top upregulated cluster DEGs. Genes in red are selected by adjusted *p* < 0.0001 (Wilcoxon rank-sum test) and |log₂FC| > 2 (See Table S1 for more details). **(h)** Time-resolved UMAP plots of glial cell types from Glia-hTDP43 flies across disease progression (1, 7, and 14 days). The top row tracks the distribution of individual glial subclusters (G*, GlEn, GlCx, GISu, GlA1, GlA2). The bottom row displays the expression of *Arc1*, mapping how the transcript expression increases in specific glial subtypes over time. **(i)** Broad-scale UMAP plots of *Arc1* expression across all captured brain cells. The panel contrasts expression in Control and Glia-hTDP43 datasets across 1, 7, and 14 days, illustrating that *Arc1* activation seen in panel (h) is a highly selective event restricted to the disease-associated glial lineage.

To trace the expression of the human *TARDBP* transgene and its pathology-seeding capability, we first mapped its age-dependent distribution across conditions. Reassuringly, while *TARDBP* expression remained strictly restricted to the glial clusters during presymptomatic stage (day 1), a distinct non-glial, neuronal subpopulation prominently contained the transgene transcript by symptomatic stages (days 7 and 14) (**Figure 2b**). This suggests a glia-to-neuron spreading of *TARDBP* transcripts within the late symptomatic brain.

Having verified transgene tracking, we next evaluated the downstream consequences of the spreading pathology on the recipient neuronal cells. We observed a marked, age-dependent reduction in cholinergic and glutamatergic neuron populations (**Figure 2a and 2c**), precisely the neuronal subtypes most vulnerable in clinical ALS and FTD^25,26^. Concomitantly, a novel, transcriptionally collapsed neuronal cluster (labelled as *non-functional*) emerged from these susceptible lineages, and to a lesser extent from GABAergic neurons (**Figure 2a; Figure S3a**). Interestingly, pathway enrichment analysis revealed that this aberrant cluster shares a highly overlapping transcriptional signature between Control and Glia-hTDP43 brains, with specific degenerative and stress pathways being further enriched under hTDP-43 pathology (e.g. *Negative regulation of cellular macromolecule biosynthetic pathway* or *Negative regulation of translation*; **Figure S3b**). Rather than inducing an entirely *de novo* artificial cell state, glial hTDP-43 expression dynamically shifts cellular proportions, triggering a massive pathological expansion of this latent aberrant neuronal population and molecularly reflecting the severe neurodegenerative cascade.

To uncover the upstream mechanisms driving this intercellular propagation, we redirected our focus to the donor glial compartment. Clustering analysis revealed the progressive emergence of a disease-specific cellular state, termed *hTDP43 Glia*, unique to the pathological Glia-hTDP43 brains (**Figure 2a**). After isolating and subclustering glial populations we observed a collocalization of other glial types, such as *Astrocyte-like glia*, with *hTDP43 Glia* (**Figure 2d**). Assuming a possible temporal differentiation process, we performed PHATE dimensionality reduction, designed to uncover global patterns from the high-dimensional topology, and combined it with trajectory inference using *Slingshot* (**Figure 2e**). Clustering on the PHATE space yielded 11 populations divided into 3 branches (A, B, C) which merged onto a shared end (ABC). This observation was supported by *Slingshot*, which produced 3 lineages aligning with the branches in the PHATE projection. Characterization of PHATE clusters showed a clear correspondence to each glial subtype (A - *Astrocyte-like glia 2*, B - *Astrocyte-like glia 1*, C - *Ensheathing glia*), with a progression from Control to Glia-hTDP43 populations alongside clusters from each branch, converging onto Glia-hTDP43-only populations (**Figure 2f**). The PAGA graph correlates with PHATE projection and Slingshot trajectory inference, providing further evidence to suggest that these existing glial subtypes undergo profound changes in response to hTDP-43 pathology, ultimately contributing to the disease state.

We next aimed to identify molecular drivers potentially implicated in this pathological glial transformation. Gene expression levels along the pseudotemporal trajectories of both *Astrocyte-like glia* and *Ensheathing glia* clusters identified *Arc1* as the most prominent molecular signature defining this transition (**Figure S4a and S4b**). Further supporting this observation, differentially expressed genes (DEGs) between glial cells from Control and Glia-hTDP43 brains highlighted *Arc1* as a top hit, exhibiting both high fold change and statistical significance of differential expression between *hTDP43 Glia* and the rest of glial types (**Figure 2g**; **Table S1**). As shown in **Figure 2h**, *Arc1* expression is almost exclusively expressed in the Glia-hTDP43 populations derived from *Astrocyte-like glia* and, to a lesser extent, from *Ensheathing glia* at presyntomatic stage (day1). Crucially, the massive *Arc1* upregulation was remarkably conserved in the Glia-hFUS cross-validation model, where its robust expression within both glial populations was already evident at day 1 (**Figure S2e**). Finally, UMAP visualization of *Arc1* expression across all cell types further demonstrates a marked age-dependent increase, peaking at the late symptomatic stage (day 14 in the Glia-hTDP43, **Figure 2i** and day 1 in Glia-hFUS, **Figure S2f**). Notably, the induction of *Arc1* expression was aligned with the emergence of functional impairments in the disease model, highlighting *Arc1* as a key gene potentially involved in ALS/FTD pathology. Given that *Arc1* retains the ancestral ability to assemble retroviral-like capsids for intercellular transport, its massive, pathology-induced overexpression in donor glia nominated it as a prime mechanistic candidate for driving the transcellular propagation of the disease. This raised the critical question of whether targeted inhibition of *Arc1*, and specifically its capsid assembly machinery, could intercept and block the intercellular transmission of TDP-43.

### 3. Genetic ablation of glial Arc1 halts non-cell-autonomous TDP-43 propagation and rescues neurodegeneration

To validate the transcriptomic increase of *Arc1*, we performed qPCR analysis in the Glia-hTDP43 model. This confirmed a highly significant upregulation of *Arc1* from day 7 onwards (**Figure 3a**). Furthermore, equivalent increase in *Arc1* was observed in the brains of the Glia-hFUS model at day 1 post-eclosion (**Figure S2g**), and in a model expressing the C9orf72-derived PR dipeptide repeat protein in glia (Glia-C9orf72 model) at day 7 post-eclosion, aligned again with the emergence of functional impairments (**Figure S2h**). This further confirms that *Arc1* elevation appears to be a convergent, universally conserved response across multiple ALS/FTD pathological contexts, establishing its broad relevance. Given that *Arc1* originated from the *gag* gene of a Ty3/Gypsy retrotransposon^27^, and retains the intrinsic capability to assemble 30 nm viral-like particles (VLPs) for intercellular cargo transfer^28,29^, we hypothesized that the massive upregulation of *Arc1* would manifest ultrastructurally in our *in vivo* fly model.

**Figure 3.**
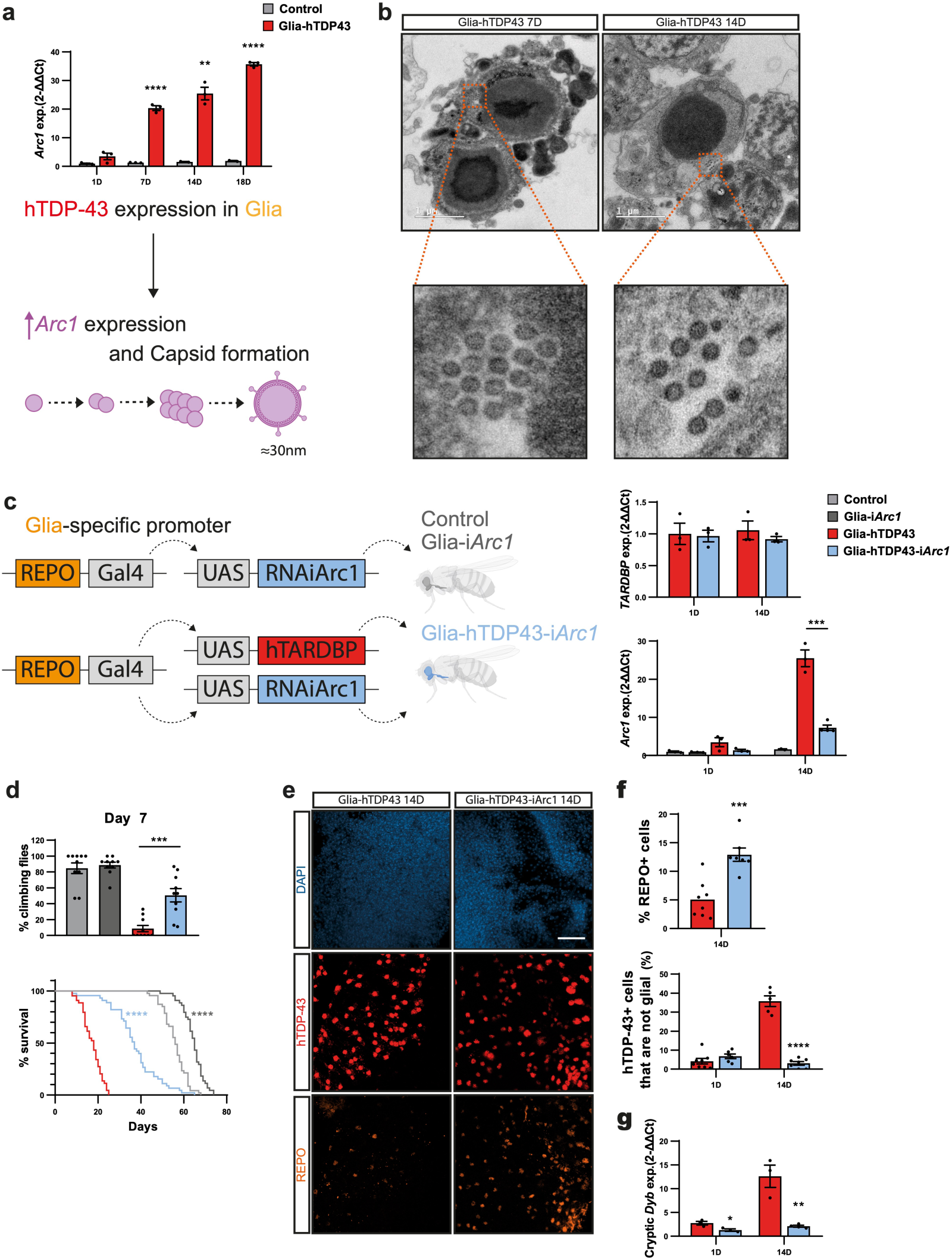
Glial *Arc1* knockdown suppresses non-cell-autonomous hTDP-43 spreading, preserving neuronal functional integrity and rescuing behavioral defects *in vivo*. **(a)** Longitudinal qPCR analysis of *Arc1* transcript expression dynamics in brains of Control and Glia-hTDP43 flies across disease progression at 1, 7, 14, and 18 days post-eclosion (top). The lower schematic outlines the working model: glial expression of hTDP-43 triggers a massive, time-dependent upregulation of *Arc1*, driving the assembly of retroviral-like capsids (VLPs 30 nm). **(b)** TEM micrographs of Glia-hTDP43 brain sections at 7 and 14 days of age. Pathological cells (top) display a massive, highly electron-dense central core structure reminiscent of severe proteinopathic inclusion bodies (putative macromolecular TDP-43 aggregates). Scale bars, 1 μm. High-magnification insets (bottom) reveal arrays of geometrically uniform, spherical VLPs matching Arc1 capsid morphology clustering within the adjacent cytoplasm. **(c)** Genetic configuration of the tissue-specific rescue system (left). The REPO-Gal4 driver was utilized to target RNA interference against *Arc1* (UAS-RNAiArc1) either in isolation (Glia-i*Arc1*) or concurrently with the human *TARDBP* transgene (Glia-hTDP43-i*Arc1*). Right panels show qPCR validation of *TARDBP* (top) and *Arc1* (bottom) expression in fly heads at days 1 and 14, confirming that *Arc1* silencing successfully blunts its pathology-induced escalation without altering primary *TARDBP* transgene transcription. **(d)** Evaluation of systemic physiological rescue. Upper histogram tracks locomotor performance via a climbing assay at day 7. Lower Kaplan-Meier lifespan survival curves map the four experimental cohorts, illustrating a robust and significant reversal of early mortality and behavioral decay upon glial *Arc1* depletion. **(e)** Confocal projections of 14-day-old adult brains from Glia-hTDP43 and Glia-hTDP43-i*Arc1* flies stained with DAPI (blue), anti–hTDP-43 (red), and anti-REPO (glia, orange). Scale bar 20 μm. **(f)** Morphometric quantifications (right) display the total percentage of REPO⁺ glial cells and the proportion of hTDP-43⁺ cells detected outside the glial boundaries (REPO⁻), demonstrating that knocking down *Arc1* halts the non-cell-autonomous escape of toxic protein seeds into neighboring tissue. **(g)** Temporal qPCR quantification of the neuron-specific *Dyb* cryptic exon expression at day 1 and day 14. Depleting glial *Arc1* significantly prevents the pathological emergence of *Dyb* cryptic splicing, demonstrating that halting capsid-mediated propagation rescues downstream nuclear loss-of-function phenotypes in recipient neurons. *Data are presented as mean ± SEM (n = 3–6 biological replicates per cohort, as detailed in individual panels). *p < 0.05, p < 0.01, ***p < 0.001, ***p < 0.0001 by Student’s t-test, one-way ANOVA with Tukey’s post-hoc test, or Mantel-Cox log-rank test for survival assays.

Strikingly, transmission electron microscopy (TEM) of Glia-hTDP43 brains revealed the accumulation of 30 nm capsid-like structures within some cells, morphologically resembling VLPs, uniquely at day 7 and 14 brain samples (**Figure 3b**). These VLPs were absent in control samples, further supporting the hypothesis of being formed from oligomerized Arc1.

To establish a causal role for Arc1/ARC in the glia-to-neuron propagation of hTDP-43, we evaluated whether its targeted depletion could halt disease progression, thereby validating this machinery as a potential therapeutic target.

We generated a *Drosophila* model combining glia-specific expression of hTDP-43 with RNAi-mediated knockdown of *Arc1* (Glia-hTDP43-i*Arc1*), alongside a control model featuring isolated glial *Arc1* knockdown (Glia-i*Arc1*). **(Figure 3c**). Consistent with our hypothesis, the Glia-hTDP43-i*Arc1* model showed a significant improvement in both motor performance and survival (**Figure 3d**). To elucidate if Glia-hTDP43-i*Arc1*-based phenotypic rescue was driven by a mechanistic blockade of intercellular glia-to-neuron transfer, we examined the spatial distribution of hTDP-43 in the brains (**Figure 3e**) and found that knockdown of *Arc1* restored the percentage of REPO-positive cells at day 14, suggesting a protective effect on glial cell survival (**Figure 3f**). Furthermore, knockdown of *Arc1* reduced the granular localization of hTDP-43 within glial processes and confined the protein to the donor glia, effectively preventing the spread of hTDP-43 to non-glial recipient cells (**Figure 3f**). To confirm the functional arrest of this toxic spreading, we assessed by qPCR the *Dyb* cryptic exon indicative of the neuronal Tbph loss-of-function. As shown in **Figure 3g**, the *Dyb* cryptic exon was not detected in the Glia-hTDP43-i*Arc1* model at either day 1 day or 14. Taken together, these findings support that Arc1 is required for packaging and transmitting competent hTDP-43 seeds to neighboring neuronal cells *in vivo*.

### 4. Stress-induced human ARC expression drives transcellular TDP-43 transmission in vitro

To examine whether the potential Arc1-mediated pathological mechanism is translatable to humans, we investigated its human ortholog, ARC, using a U2OS cell line with inducible TDP-43-GFP expression. Under basal conditions, *ARC* expression is minimal; however, upon exposure to oxidative and ribotoxic stress via sodium arsenite (NaAsO₂) to trigger TDP-43 mislocalization, *ARC* mRNA levels were robustly upregulated (**Figure 4a**). Broadening the physiological relevance of this response, we found that *ARC* is also highly upregulated in U2OS cells following the isolated overexpression of ALS/FTD-linked factors (TDP-43, C9orf72-DPRs, or hnRNPA2) or exposure to the environmental neurotoxin β-Methylamino-L-alanine (BMAA) (**Figure S5**). This demonstrates that *ARC* induction a convergent molecular signature across diverse proteotoxic insults related to FTD/ALS. Most intriguingly, imaging analysis revealed that NaAsO₂ treatment not only triggered the cytoplasmic coalescence of TDP-43 but also drove ARC to form granule-like structures that colocalized with TDP-43 (**Figure 4b**).

**Figure 4.**
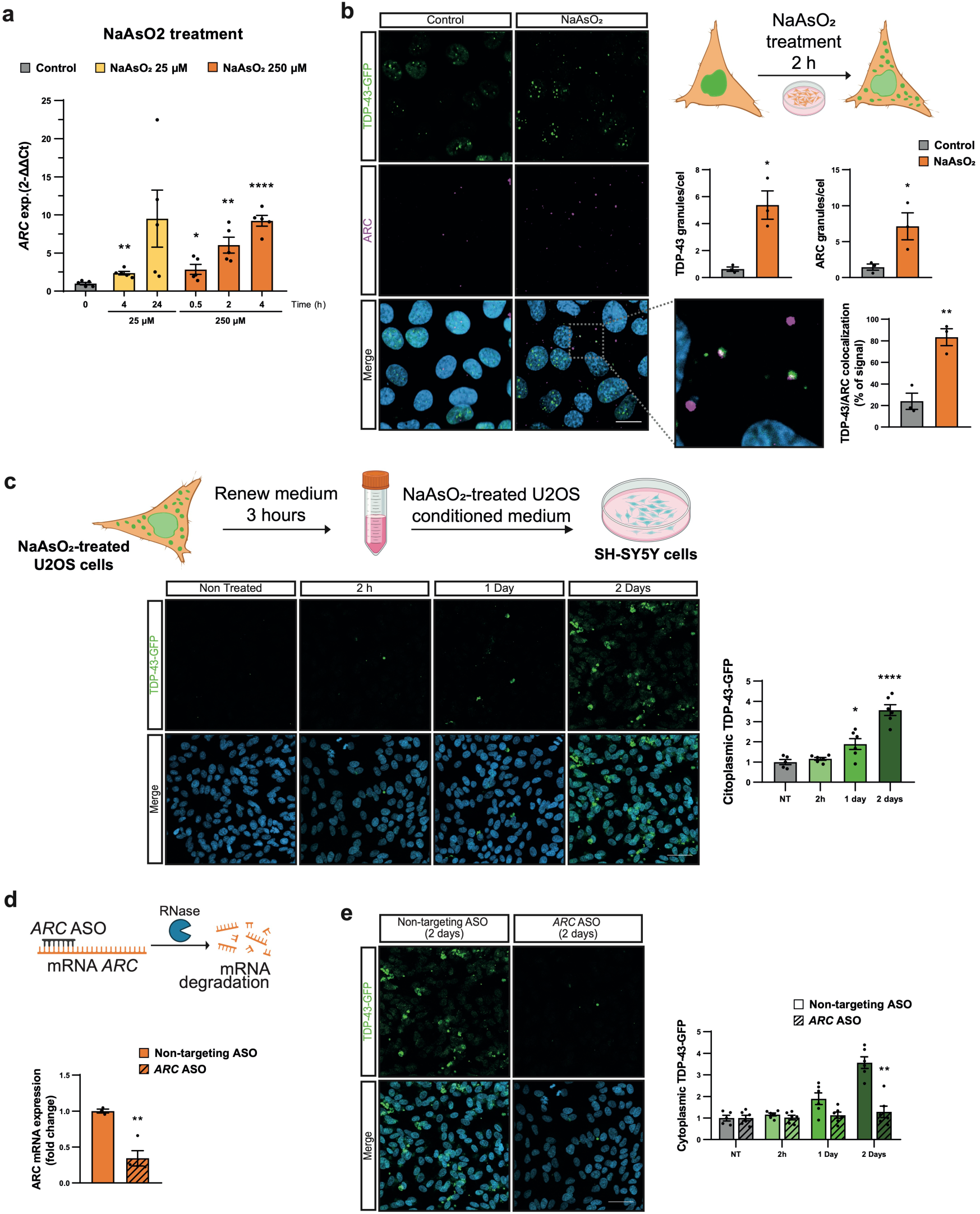
Arsenite-induced human ARC expression drives TDP-43 incorporation into the secretory phenotype and mediates intercellular seed transmission *in vitro*. **(a)** Dose- and time-dependent qPCR profiling of human *ARC* mRNA expression in U2OS cells exposed to NaAsO_2_ at concentrations of 25 and 250 µM. **(b)** Representative confocal micrographs of donor cells expressing TDP-43-GFP (green) and immunostained for endogenous ARC (magenta) under baseline control conditions or following a 2 hours acute NaAsO_2_ stress window. Upper schematic outlines the experimental design. Histograms clarify the morphometric properties of the resulting stress granules, quantifying (i) TDP-43 granules per cell, (ii) ARC granules per cell, and (iii) the exact percentage of physical TDP-43/ARC signal colocalization within cytoplasmic condensates. **(c)** Schematic representation of the conditioned medium transfer workflow (top). Stressed donor cells undergo a thorough washing protocol, medium is renewed for 3 hours to harvest the natively secreted cargo, and this conditioned medium is subsequently applied to recipient SH-SY5Y neuroblastoma cells. Longitudinal confocal panels capture the real-time uptake of cytoplasmic TDP-43-GFP inside recipient cells across Non-Treated (NT), 2 hours, 1 day, and 2 days timepoints. The adjacent graph tracks the progressive escalation of internalized cytoplasmic TDP-43-GFP fluorescence. **(d)** Operational diagram of the gapmer antisense oligonucleotide strategy targeting *ARC* transcripts (top). The lower bar chart validates target knockdown efficacy via qPCR, demonstrating a significant reduction in total *ARC* mRNA levels in the treated cells. **(e)** Confocal micrographs of recipient SH-SY5Y cells at day 2 post-incubation with conditioned medium derived from either non-targeting ASO or *ARC* ASO-silenced donor cells. The temporal bar graph tracks recipient cytoplasmic TDP-43-GFP signal accumulation across dynamic intervals (NT, 2 hours, 1 day, and 2 days), demonstrating that depleting donor *ARC* expression effectively disrupts the transcellular transmission and propagation of pathological seeds. *All representative fluorescence images were captured under identical optical constraints, lasers, and magnification parameters. Data are plotted as mean ± SEM (n = 3–4 independent biological replicates). *p < 0.05, p < 0.01, ***p < 0.001, \*\*\**p < 0.0001* by Student’s t-test or two-way ANOVA followed by Tukey’s post-hoc test for the propagation kinetics.

Given that human *ARC* expression is induced by stress and physically colocalizes within cytoplasmic TDP-43 granules, we next investigated whether this machinery has the capacity to transfer TDP-43 to naïve cells. For this purpose, we established a human cell-based TDP-43-GFP transfer assay. U2OS donor cells were treated with 250 µM NaAsO₂for 2 hours, a paradigm identified in **Figure 4a** as the optimal condition for optimal *ARC* induction. Following a 3-hour washout and recovery period to ensure toxin clearance and allow sufficient time for vesicular cargo accumulation, the conditioned medium was harvested and transferred to wild-type SH-SY5Y recipient cells. Quantitative analysis showed a time-dependent increase in the internalization of TDP-43-GFP within SH-SY5Y cells (**Figure 4c**), demonstrating that factors secreted into the extracellular space by the donor cells are sufficient to drive the intercellular transfer of pathological TDP-43 in human cells.

We next examined the involvement of ARC in the intercellular transmission of TDP-43 in the U2OS model. To this end, we utilized our human *in vitro* transfer assay, employing a targeted gapmer antisense oligonucleotide (ASO) to efficiently knockdown *ARC* expression in donor U2OS TDP-43-GFP cells prior to NaAsO_2_ stress (**Figure 4d**). Remarkably, *ARC* silencing in the donor cells diminished the presence of pathogenic factors in the conditioned medium, resulting in a marked reduction of TDP-43-GFP internalization into the naïve recipient SH-SY5Y cells (**Figure 4e**).

Collectively, these findings show that the model initially derived from the scRNAseq fly data translates into human cells, establishing ARC as a targetable protein required for transcellular propagation of TDP-43 proteinopathy.

### 5. Computational modeling identifies Lenacapavir as a preferential ligand of the ARC/Arc1 capsid-forming interface

Having established that Arc1/ARC retroviral-like machinery is required for TDP-43 propagation, we next asked whether this process could be pharmacologically intercepted at the level of capsid-like assembly. Because the Gag domain of ARC/Arc1 drives virus-like particle formation, we reasoned that small molecules targeting retroviral capsid interfaces might also engage the structurally related ARC/Arc1 assembly interface. Lenacapavir, an FDA-approved HIV-1 capsid forming modulator, binds the HIV-1 capsid at an inter-subunit pocket formed between neighboring capsid subunits (**Figure S6**), ultimately leading to aberrant capsid formation^30^. We therefore tested whether Lenacapavir could occupy an analogous interface in *Drosophila* Arc1 and human ARC.

To avoid relying on docking alone, we implemented a stepwise computational pipeline integrating: structural modeling, molecular dynamics (MD), induced-fit docking (IFD), binding-pose metadynamics (BPMD) and absolute binding free-energy (ABFE) calculations. For *Drosophila* Arc1, we used the experimentally resolved hexameric capsid assembly (PDB: 6TAS) and extracted the minimal dimeric interface containing the predicted Lenacapavir-binding region (**Figure 5a**). For human ARC, where no experimentally resolved hexameric structure is available, we generated a capsid-forming assembly using AlphaFold3 (**Figure 5f; Figure S7a and S7b**) and validated the stability of the predicted structure by independent MD simulations prior to ligand evaluation (**Figure S7c and S7d**).

**Figure 5.**
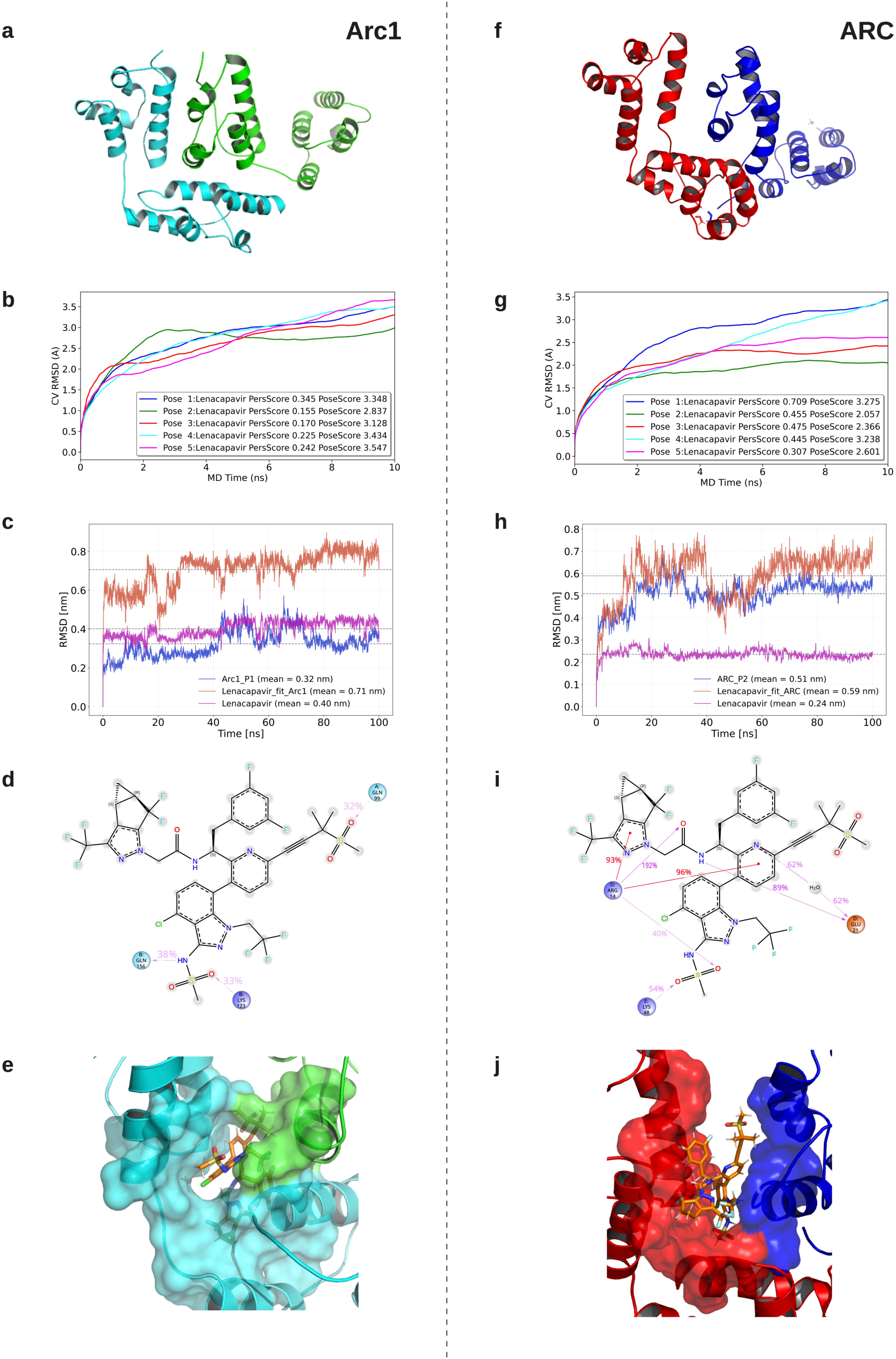
*In silico* docking and atomistic molecular dynamics simulations identify structurally conserved capsid pockets in *Drosophila* Arc1 and human ARC as stable targets for Lenacapavir. **(a)** PDB-derived 3D structural model of the *Drosophila* Arc1 domain mapping the spatial topography of the target binding pocket (Pocket 1, P1) selected for small-molecule virtual screening. **(b)** Conformational convergence profiling illustrating the cluster-vector Root-Mean-Square Deviation (CV RMSD, in Å) over a 10 ns molecular dynamics (MD) refinement window for the top five predicted binding poses of Lenacapavir within the Arc1 P1 cavity. Persistence and pose scores are detailed for each candidate trajectory. **(c)** Structural stability assessment during a prolonged 100 ns production-phase all-atom MD simulation. Trajectory plots track the structural drift (RMSD, in nm) of the standalone Arc1 pocket backbone, the fitted ligand-protein complex, and the bound Lenacapavir ligand over time. **(d)** 2D ligand-protein interaction topology detailing the specific amino acid coordinates, hydrogen bonds, and hydrophobic or electrostatic contact networks stabilizing the Lenacapavir molecule inside the *Drosophila* Arc1 binding pocket. **(e)** 3D solvent-accessible surface rendering of the *Drosophila* Arc1 pocket (cyan/green overlay), showcasing the geometric accommodation and volumetric fit of Lenacapavir (orange sticks) inside the core cavity. **(f)** AlphaFold-derived 3D structural model of the human ARC capsid domain, highlighting the equivalent conserved binding pocket (Pocket 2, P2) evaluated for orthologous drug targeting. **(g)** CV RMSD trajectory curves over a 10 ns structural refinement scale for the top five independent docking poses of Lenacapavir within the human ARC P2 pocket. **(h)** 100 ns productionall-atom MD simulation tracking the trajectory stability (RMSD, in nm) of the human ARC P2 pocket backbone, the fitted ligand complex, and the free Lenacapavir molecule inside the human protein environment. **(i)** 2D chemical interaction map resolving the coordinating amino acid residue network and intermolecular forces stabilizing Lenacapavir within the human ARC P2 binding domain. **(j)** 3D conformational surface map displaying Lenacapavir (yellow sticks) deeply wedged into the inter-domain structural pocket interface formed between coordinating helical bundles (surface rendered in red and blue) of the human ARC architecture. *All atomistic molecular dynamics trajectories were simulated under explicit solvent conditions (150 mM KCl) at a constant physiological temperature (298.15 K). Structural models and surface potentials were resolved utilizing AMBER ff14SB force fields and energy minimized prior to production phase tracking.

In both systems, candidate ligand poses were first identified by targeted docking at the inter-subunit cavity and subsequently clustered to remove redundant binding modes. The resulting docking poses were subsequently refined by BPMD to assess their dynamic persistence and identify the most robust binding modes. The best-ranked poses showed stable behavior during BPMD and were therefore prioritized for further analysis, although they remained outside the stringent Clark et al. (2016)^31^ criteria for fully stable complexes (PoseScore ≤ 2.0 Å and PersScore ≥ 0.6). Accordingly, BPMD was not interpreted as definitive evidence of binding stability, but rather as an initial filtering step to discard clearly unfavorable poses. Selected complexes were then subjected to extended 100-ns MD simulations, which served as a second stability filter under unbiased conditions. Poses were discarded when stabilization relied predominantly on water-mediated interactions, when the ligand exhibited large RMSD fluctuations indicative of pocket escape, or when no conformational convergence was observed during the trajectory. Finally, the remaining poses were evaluated by ABFE calculations

In *Drosophila* Arc1, Lenacapavir adopted a binding mode within the inter-monomer cavity that remained comparatively stable throughout the subsequent analyses. During MD simulations, the selected pose displayed RMSD convergence relative to the protein backbone, suggesting that the ligand reached a stable or semi-stable local minimum within the cavity (**Figure 5b and 5c; Figure S8a**). Interaction analysis further revealed that Lenacapavir remained accommodated through a network of hydrogen bonds, salt bridges, primarily involving its sulfonamide and sulfone groups and surrounding polar residues (**Figure 5d and 5e**). Although the overall interaction pattern partially resembled the binding mode observed in the HIV-1 capsid–Lenacapavir complex, interaction persistence was lower in Arc1, indicating a weaker and more dynamic stabilization network. Despite these differences, ABFE calculations predicted a favorable affinity for the Arc1–Lenacapavir complex, with ΔG = −10.07 ± 0.39 kcal/mol, corresponding to a predicted Kd of 42 nM (**Table S2**).

We next applied the same computational framework to human ARC. Following validation of the AlphaFold3-derived capsid-like assembly by MD simulations, docking and BPMD identified a putative Lenacapavir binding mode at the corresponding dimeric interface (**Figure 5f**). Subsequent 100-ns MD simulations indicated that the ligand remained stably accommodated within the pocket without perturbing the dimeric assembly (**Figure 5g and 5h; Figure S8b**). At the atomic level, the predicted binding mode preserved key polar and electrostatic contacts mediated by the sulfone/sulfonamide region, together with aromatic and cation-associated interactions that contributed to ligand stabilization (**Figure 5i and 5j),** consistent with partial conservation of the interaction pattern observed in the HIV-1 capsid complex. ABFE calculations predicted strong binding to human ARC, with ΔG = −8.67 ± 0.22 kcal/mol and an estimated Kd of approximately 0.44 µM (**Table S2**). Although the absolute affinity should be interpreted with caution because the human ARC assembly is model-derived, these results indicate that the human ARC interface can accommodate Lenacapavir in a stable and energetically favorable binding mode.

To test whether this effect reflected general capsid-inhibitor chemistry or a more specific structural complementarity, we performed the same analysis with Bersacapavir, a hepatitis B virus (HBV) capsid assembly modulator used here as a negative control. Despite sharing a sulfonamide-containing scaffold, Bersacapavir failed to establish comparably stable binding modes at either the Arc1 or human ARC interfaces. In both systems, predicted interactions were weaker, less persistent, and more frequently water-mediated (**Figure S8c and S8d**). In particular, reduced conformational stability was observed in the human ARC pocket relative to Lenacapavir. Consistent with these findings, ABFE calculations predicted substantially weaker affinities for Bersacapavir, with estimated Kd values of approximately 21–41 µM for Arc1 and approximately 576 µM for human ARC, representing a staggering 1,000- to 10,000-fold decrease in predicted binding potency relative to Lenacapavir (**Table S2**).

Together, these *in silico* data identify Lenacapavir as a preferential ARC/Arc1 capsid-interface ligand and provide a molecular rationale for its selective activity relative to Bersacapavir. The results support a model in which the three-dimensional architecture of Lenacapavir, rather than the presence of a sulfonamide group alone, enables productive engagement of the ARC/Arc1 inter-subunit cavity, thereby nominating Lenacapavir as a candidate pharmacological modulator of ARC-dependent TDP-43 propagation.

### 6. Pharmacological modulation of ARC/Arc1 capsid assembly with Lenacapavir arrests TDP-43 propagation and rescues neurodegenerative phenotypes

Encouraged by our *in silico* structural predictions, we evaluated the therapeutic of Lenacapavir *in vivo* using the Glia-hTDP43 *Drosophila* model. Strikingly, dietary administration of Lenacapavir (at 0.5 µM and 2.5 µM) significantly improved motor activity and extended overall survival **(Figure 6a)**. This pharmacological rescue closely resembled the phenotypic rescue achieved via genetic *Arc1* knockdown, validating the capsid interface as an actionable *in vivo* target. Further demonstrating its broad translational potential, Lenacapavir treatment (2.5 µM) also prolonged lifespan in the Glia-hFUS model **(Figure 6b)**, confirming its efficacy across distinct FTD/ALS proteinopathies.

**Figure 6.**
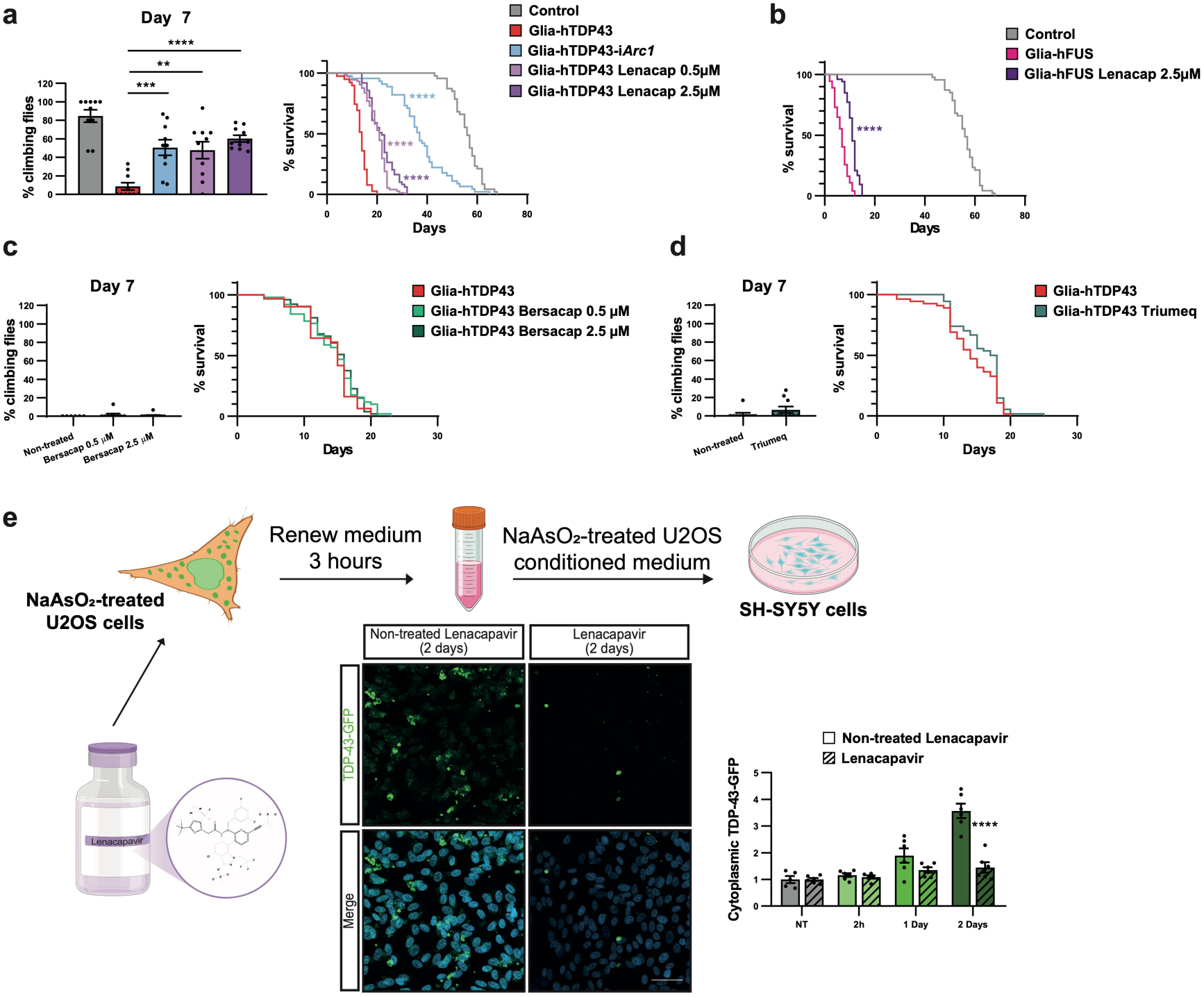
Pharmacological inhibition of capsid assembly by Lenacapavir selectively mitigates non-cell-autonomous TDP-43 neuropathology and behavioral decay. **(a)** Motor performance evaluated via negative geotaxis assays at day 7 (left) and longitudinal Kaplan-Meier lifespan survival curves (right) of Glia-hTDP43 flies treated with Lenacapavir (0.5 and 2.5 µM). **(b)** Lifespan survival curves of a humanized glial FUS proteinopathy model (Glia-hFUS) treated with Lenacapavir (2.5 µM), confirming the disease-specific nature of the capsid-targeted intervention. **(c)** Locomotor performance at day 7 (left) and survival curves (right) of Glia-hTDP43 flies treated with Bersacapavir (0.5 and 2.5 µM), acting as a structurally inactive chemical analogue control to confirm structure-activity relationship (SAR) specificity. **(d)** Locomotor performance (left) and lifespan curves (right) of Glia-hTDP43 flies treated with the clinical multi-class antiretroviral regimen Triumeq, illustrating the lack of therapeutic efficacy when targeting canonical reverse transcriptase and integrase enzymes rather than physical capsid assembly. **(e)** Schematic representation of the *in vitro* propagation blockade assay (top). Conditioned medium harvested from NaAsO_2_-stressed donor U2OS cells treated with or without Lenacapavir was transferred to recipient SH-SY5Y neuroblastoma cells. Representative confocal micrographs (bottom left) and longitudinal automated morphometric tracking (bottom right) quantify recipient cytoplasmic TDP-43-GFP accumulation across dynamic intervals (NT, 2 hours, 1 day, and 2 days), validating that small-molecule capsid disruption potently halts intercellular seed transmission. Scale bar = 10 μm. Data shown as mean ± SEM (n = 3–6 independent biological replicates or cohorts, as detailed in individual panels). Statistical significance: \**p* < 0.05, **p < 0.01, \*\*\**p* < 0.001, ****p < 0.0001 by log-rank Mantel-cox test for the longevity assays, or two-way ANOVA followed by Tukey’s post-hoc test for the Lenacapavir propagation assay).

To empirically validate the structural specificity predicted by our thermodynamic models, we treated the Glia-hTDP43 flies with Bersacapavir. As a HBV capsid assembly modulator, Bersacapavir targets viral capsids but lacks the specific stereochemical conformation required to stably bind the Arc1 dimeric interface, as demonstrated by our ABFE calculations. Consistent with these *in silico* predictions, Bersacapavir administration (at 0.5 µM and 2.5 µM) failed to provide any protective effect on motor performance or survival (**Figure 6c**). This negative control treatment confirms that specific structural matches to ARC/Arc1 binding sites are required to adequately intercept the spread of hTDP-43.

We next sought to rule out the possibility that TDP-43 propagation is mediated by the generalized activation of fully competent endogenous retroviruses (ERVs) or other canonical retrotransposons. To this end, we employed Triumeq as another mechanistic negative control *in vivo*. Triumeq is a frontline combination antiretroviral therapy comprising an integrase inhibitor (Dolutegravir) and two nucleoside reverse transcriptase inhibitors (Abacavir and Lamivudine). Unlike active ERVs, which critically rely on reverse transcriptase and integrase enzymes to execute their pathogenic lifecycles, the domesticated *ARC/Arc1* gene encodes exclusively a Gag-like capsid domain. We reasoned that if the intercellular transfer was driven by classical retrotransposons, Triumeq should confer phenotypic protection. Conversely, if propagation depends strictly on the reverse transcriptase/integrase-independent capsid assembly of Arc1, this drug cocktail would be therapeutically inefficient. As anticipated, the Glia-hTDP-43 model treated with Triumeq showed no significant differences in motor activity or survival compared to the same untreated model (**Figure 6d**). Therefore, the Gag-like domain specific from the ARC/Arc1 protein is the required element to target vehicle-driven TDP-43 spreading.

To unequivocally link this *in vivo* phenotypic rescue to the specific blockade of transcellular seeding, we evaluated Lenacapavir in our human cellular propagation assay. Concordant with the *Drosophila* data, Lenacapavir treatment of the donor U2OS cells significantly reduced the release of pathogenic cargo, resulting in a near-complete suppression of intercellular propagation of TDP-43-GFP internalization in naïve SH-SY5Y recipient cells (**Figure 6e**). This directly validates that Lenacapavir intercepts the physical propagation of TDP-43 mediated by human ARC.

Collectively, these findings provide compelling evidence that ARC/Arc1 VLPs are indispensable vehicles for the intercellular propagation of TDP-43 proteinopathy. Furthermore, our results points out that repurposing FDA-approved retroviral capsid inhibitors, such as Lenacapavir, constitutes a highly specific, immediately translatable strategy to arrest the anatomical spread of neurodegeneration.

## DISCUSSION

The progressive anatomical spread of neurodegeneration defines both ALS and FTD. Clinically, this remains an untreatable feature. In this study, we elucidate the structural and mechanistic basis driving this transcellular cascade. We demonstrate that ARC/Arc1, a domesticated retrotransposon-derived protein classically restricted to homeostatic synaptic plasticity^32,33^, is pathologically repurposed. Under severe proteotoxic stress, ARC/Arc1 shifts its function from synaptic formation to assemble retroviral-like VLPs that actively package and transmit toxic TDP-43 seeds to recipient cells.

*In vitro* models have thoroughly mapped the prion-like, cell-to-cell transmission of misfolded TDP-43^11,34,35^. However, tracking this dynamic dissemination *in vivo* has been difficult to prove univocally. For instance, existing animal paradigms rely heavily on the stereotaxic injection of patient-derived brain extracts^36^, which introduces critical confounding variables like additional proteins mediating the transmission of TDP-43 or acting through additional pathways. Localized mechanical tissue injury, blood-brain barrier disruption, and the non-physiological delivery of exogenous material act as complementary ways to exacerbate ALS pathology but do not directly prove the transmission of misfolded TDP-43 as the primary approach. In this study we bypassed these limitations by developing a genetic model that strictly confines conditional human TDP-43 expression to glia in *Drosophila melanogaster*. This model provides the spatiotemporal control needed to trace directional pathology transmission from a precise cellular origin to recipient neurons inside an intact CNS. Crucially, our model proves that pathologically stressed glia export functional seeds via endogenous pathways. This establishes an ideal *in vivo* platform to genetically and pharmacologically dissect the transmission machinery.

Recent landmark studies highlight a critical mechanistic coupling. Internalized pathogenic TDP-43 aggregates readily seed the misfolding of endogenous TDP-43, directly linking cytoplasmic accumulation with subsequent nuclear depletion^7,8^. We demonstrate that ARC VLPs act as the physiological delivery vehicles driving this mechanism. By exporting toxic seeds, ARC capsids trigger a dual pathological hit. They first accelerate the cytoplasmic gain-of-function (aggregate accumulation) and then drive the neuronal nuclear loss-of-function (evidenced by the neuronal *Dyb* cryptic exon)^24^. This effectively unifies the two core pathophysiological features of TDP-43 pathology.

While the exact biochemical rules dictating how ARC would segregate, package, and export these functions are not fully mapped, our findings provide a compelling mechanistic framework for this process. Physiologically, ARC VLP assembly requires association with mRNA or specific structural RNA motifs^37^. Under the pathological conditions of ALS and FTD evaluated here, ARC redistributes and localizes to stress granules (SGs) following ribotoxic stress. This behavior perfectly matches its molecular architecture. The C-terminal motif of ARC features a prominent intrinsically disordered region^38^ that drives liquid-liquid phase separation and partitioning into dynamic SG condensates. We hypothesize this biophysical environment serves as the primary hub for capsid assembly. Furthermore, recent biochemical profiling reveals that mammalian ARC possesses potent nucleic acid chaperone activity. It facilitates RNA strand exchange and actively remodels local RNA structures^37^. Within the dense, multivalent RNA-protein network of a SG, ARC likely does not form a passive structural shell. Instead, it may actively entangle, unfold, and encapsidate non-cognate, stress-associated transcripts, such as *TARDBP* mRNA, prior to vesicular export. Alternatively, capsid assembly within SGs might simply co-package adjacent, aggregate-prone ribonucleoproteins into nascent extracellular vesicles (EVs), since ARC lacks the sequence-specificity domain that would restrict capsid formation to its own RNA. In either scenario, the outcome is identical. ARC VLPs function as structural shuttles that export the surrounding stress machinery, including TDP-43 seeds. Consequently, genetically or pharmacologically disrupting ARC assembly blocks this vesicular loading. Toxic seeds are retained within the donor cell, terminating the propagation cascade. While this intracellular containment may exacerbate toxicity for the isolated donor cell under chronic stress, it successfully protects the network. The functional integrity of neighboring healthy cells is preserved.

*ARC* is classically defined as an early expression gene restricted to homeostatic neurons^39^. However, our single-cell transcriptomic profiling identified a marked, disease-associated upregulation of the *Drosophila* ortholog specifically within the glial compartment. This proves that ARC VLP-forming machinery can be induced in non-neuronal lineages experiencing severe cellular stress. Indeed, our findings establish *ARC* as a highly sensitive responder to both proteotoxic and ribotoxic insults. Its expression is triggered by specific ALS/FTD-associated proteins and by chemical stressors like NaAsO_2_. This broad stress-responsiveness is independently corroborated. Public transcriptomic datasets^40^ consistently show *ARC* as a highly upregulated transcript following NaAsO_2_ exposure in human osteosarcoma cells. This pattern of aberrant, non-neuronal ARC expression activation under pathological strain is further supported by recent oncology literature. In malignant gliomas, upregulated ARC drives disease progression by promoting intercellular RNA transfer via VLPs^41^. Consequently, we observe robust *ARC/Arc1* induction across diverse *in vivo* and *in vitro* models (TDP-43, FUS, and C9orf72). This signals a fundamental mechanistic insight. The activation of this retroviral-like pathway represents a convergent pathological node across entirely different forms of neurodegenerative stress.

Identifying ARC as the structural vehicle for proteinopathy propagation exposes a highly specific therapeutic vulnerability. Accumulating evidence shows that endogenous retroviruses are upregulated in ALS^18,19,42^. This observation directly motivated clinical trials, such as the Lighthouse II study, to evaluate combination antiretroviral therapies like Triumeq. However, Triumeq specifically inhibits reverse transcriptase and integrase. These are essential enzymes for canonical viral replication. The domesticated *ARC* gene, conversely, consists exclusively of a Gag-like capsid domain. It lacks these enzymatic functions entirely. Therefore, these standard antiretroviral interventions are mechanistically bypassed. This molecular divergence elegantly explains the absent or modest clinical efficacy observed in recent human trials^43,44^ and in transgenic TDP-43 mouse models^45^. Our *in vivo* data empirically validates this limitation. Triumeq completely failed to modify disease progression or motor deficits in our model. In stark contrast, Lenacapavir, a targeted capsid assembly modulator, significantly blocked TDP-43 propagation and rescued functional phenotypes *in vivo* and *in vitro*. The therapeutic implication is clear. Strategies aimed at halting pathological spreading must target the structural assembly of the ARC capsid, not the enzymatic replication pathways.

Finally, emerging evidence links ARC VLPs to the propagation of Tau pathology in Alzheimer’s disease^21^. Combined with our findings in ALS and FTD, ARC capsid machinery may represent a conserved, cross-disease pathway for the transcellular spread of distinct neurodegenerative proteinopathies. Repurposing clinically approved retroviral capsid modulators, such as Lenacapavir, may offer a translational strategy to ameliorate the progression of this family of neurodegenerative disorders.

## STAR METHODS

### KEY RESOURCES TABLES

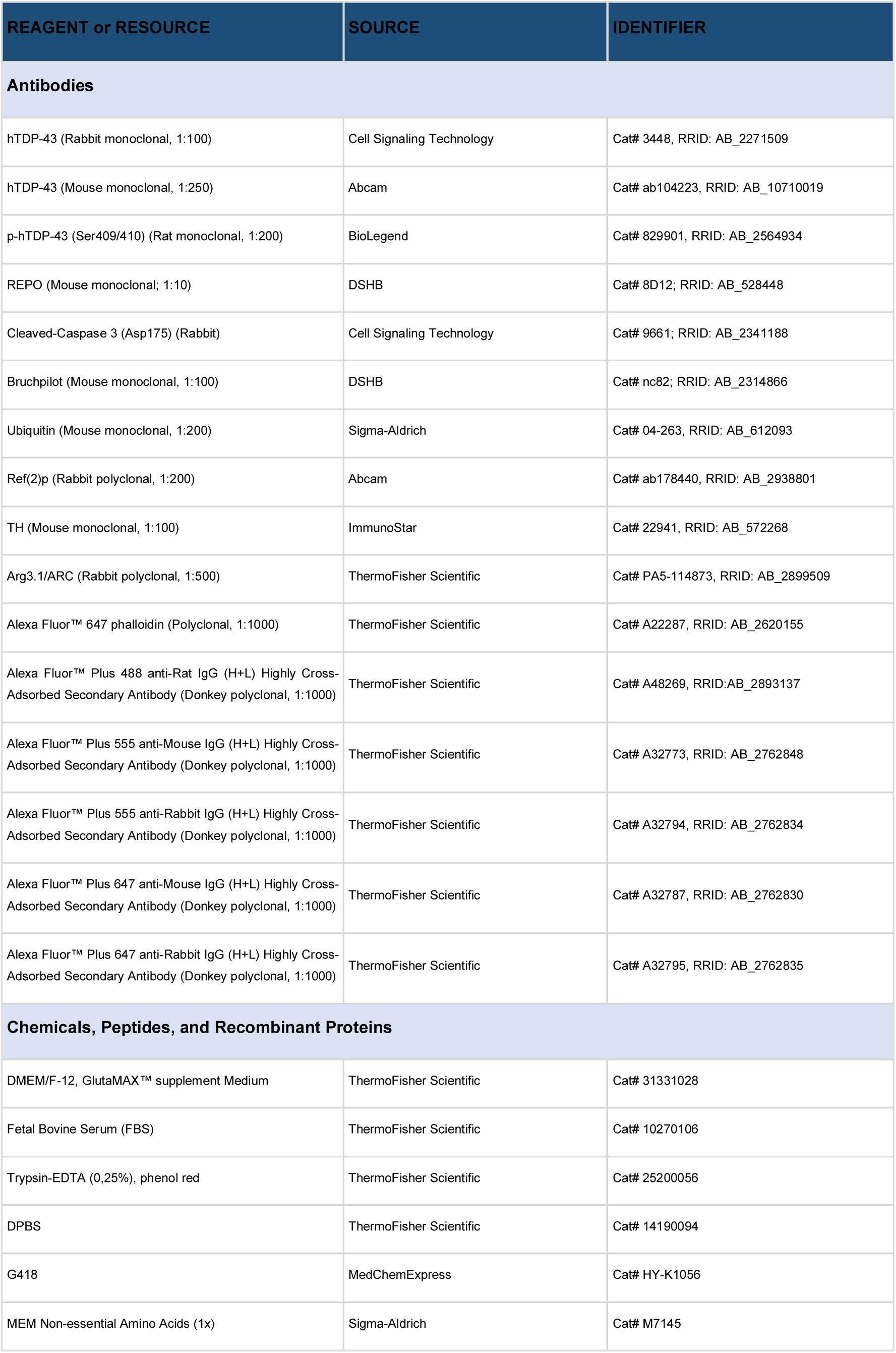
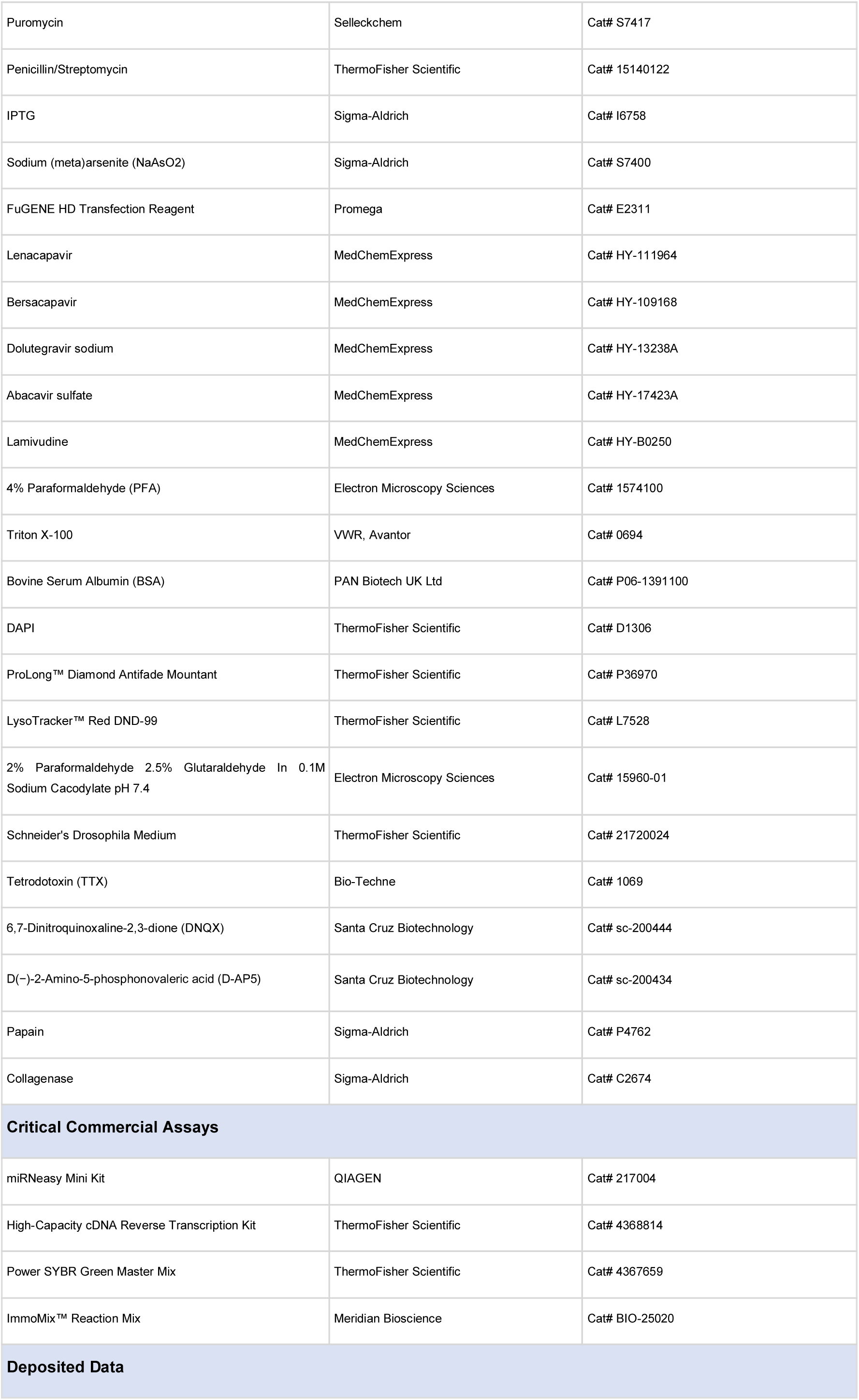
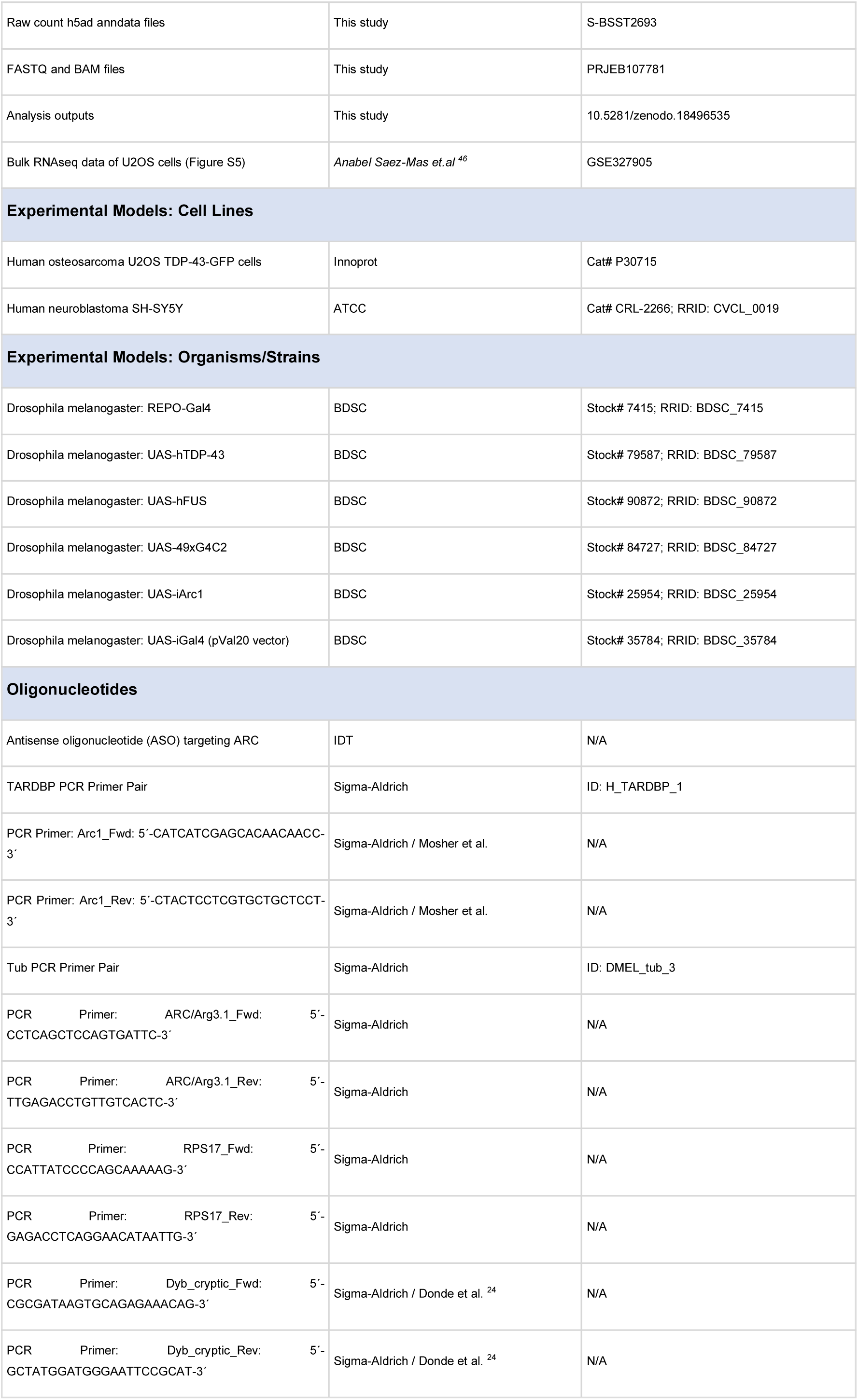
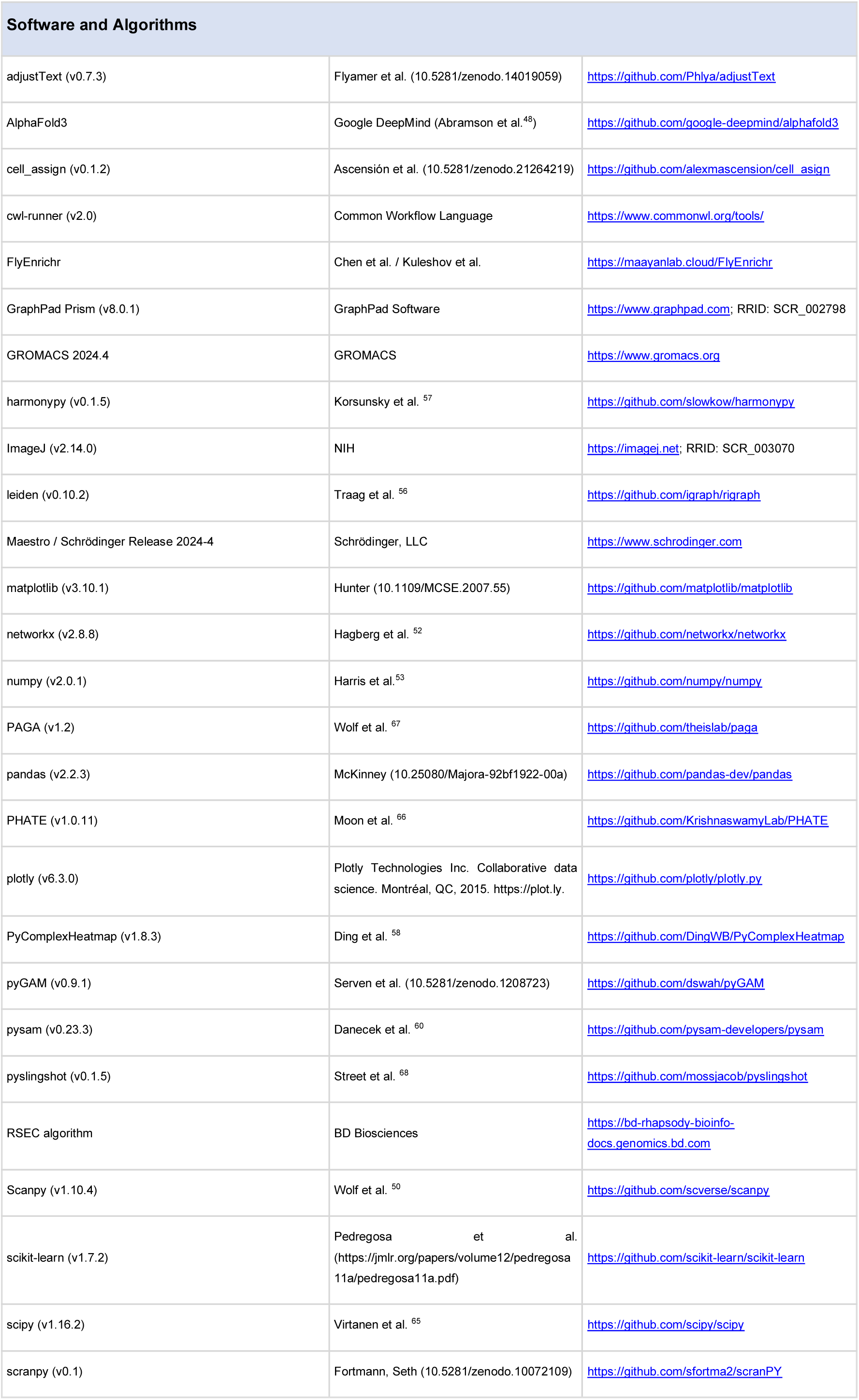
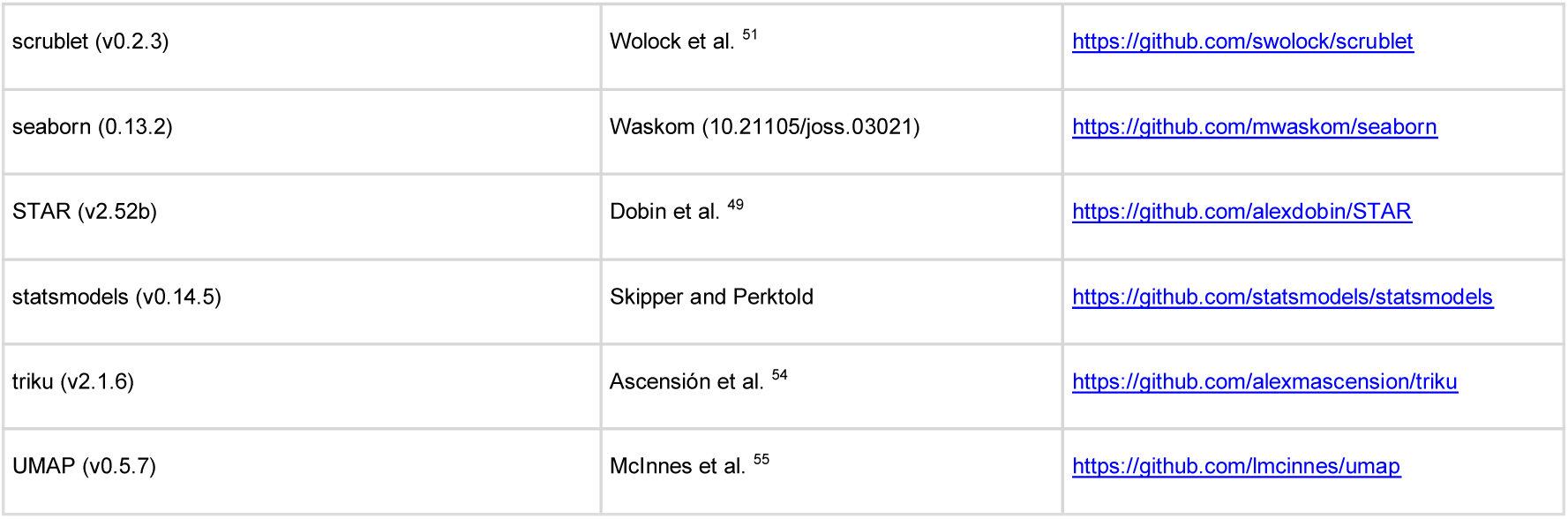

## EXPERIMENTAL MODEL AND SUBJECT DETAIL

### Cell Culture

Human osteosarcoma U2OS cells (Innoprot, Bizkaia, Spain) stably expressing GFP-tagged TDP-43 were maintained in DMEM/F-12, GlutaMAX^TM^ supplement Medium (ThermoFisher Scientific, Waltham, MA), 10 % fetal bovine serum (FBS; ThermoFisher Scientific, Waltham, MA), 300 µg/mL G418 (MedChemExpress, Monmouth Junction, NJ), 1x MEM Non-essential Amino Acids (Sigma-Aldrich, St. Louis, MO) and 3 µg/mL puromycin (Selleckchem, Houston, TX) at 37 °C in a humidified atmosphere containing 5 % CO_2_. Expression of TDP-43-GFP was induced by the addition of 1 mM IPTG (Sigma-Aldrich, St. Louis, MO) to the culture medium.

Human neuroblastoma SH-SY5Y cells (ATCC, Manassas, VA) were cultured in DMEM/F-12, GlutaMAX^TM^ supplement Medium (ThermoFisher Scientific, Waltham, MA), 10 % FBS (ThermoFisher Scientific, Waltham, MA) and 100 mg/mL penicillin/streptomycin (ThermoFisher Scientific, Waltham, MA) under the same conditions.

#### Drosophila strains

All the *Drosophila* strains were obtained from Bloomington Stock Centre (BDSC; Bloomington, IN). The driver used for the expression in glia cells was REPO (7415; RRID: BDSC_7415). This driver *Drosophila* strain was crossed with other strains carrying sequences to express hTDP-43 (79587; RRID: BDSC_79587), hFUS (90872; RRID: BDSC_90872), hC9orf72 (84727; RRID: BDSC_84727) or to knockdown *Arc1* (25954; RRID: BDSC_25954). The final experimental genotypes were Glia-hTDP43 (UAS-hTDP-43/REPO-Gal4), Glia-hFUS (UAS-hFUS/REPO-Gal4), Glia-hC9orf72 (UAS-49xG_4_C_2_/REPO-Gal4), Glia-i*Arc1* (UAS-i*Arc1*/REPO-Gal4) and Glia-hTDP43-i*Arc1* (UAS-hTDP-43-i*Arc1*/REPO-Gal4). For the control model, we crossed the driver line REPO with a strain carrying a pVal20 vector to knockdown *Gal4* (35784; RRID: BDSC_35784). The final genotype was Control (UAS-i*Gal4*/REPO-Gal4).

Stocking flies were housed at 22 °C, 70 % humidity and 12 h/12 h light/darkness cycle and experimental flies at 26 °C, 70 % humidity and 12 h/12 h light/darkness cycle.

## METHOD DETAILS

### Cell Treatments

For stress granule induction, U2OS cells stably expressing TDP-43-GFP were exposed to 250 µM sodium (meta)arsenite (NaAsO_2_) (Sigma-Aldrich, St. Louis, MO) for 2 h. For gene silencing, cells were treated 24 h with an antisense oligonucleotide (ASO) (120 µM) (IDT, Coralville, IA) targeting ARC following the manufacturer’s protocol. Control cells were processed in parallel without treatment.

As an experimental treatment, Lenacapavir (MedChemExpress, Monmouth Junction, NJ), a retroviral drug that acts as a HIV capsid modulator, was applied to U2OS cells at a final concentration of 0.1 nM for 2 h, simultaneously with NaAsO_2_ exposure. Control cells were processed in parallel under identical conditions without treatment.

#### Drosophila Treatments

All flies were collected in prepared tubes containing standard feed with or without 0.5 or 2.5 µM Lenacapavir (MedChemExpress, Monmouth Junction, NJ), 0.5 or 2.5 µM Bersacapavir (MedChemExpress, Monmouth Junction, NJ), or Triumeq, a combination of 100 nM Dolutegravir sodium (MedChemExpress, Monmouth Junction, NJ), 20 µM Abacavir sulfate (MedChemExpress, Monmouth Junction, NJ) and 1 µM Lamivudine (MedChemExpress, Monmouth Junction, NJ).

#### *Drosophila* Functional assays

For longevity tests, 50 adult female flies (five per tube) were selected from each strain. Dead flies were counted every 2 days and tubes were replaced minimum every 7 days. For locomotor activity, adult flies were tested on days 7 and 14 in groups of five (the same groups as in the longevity test). They were placed in a tube with a line drawn outside the tube at a height of 8 cm from the bottom. The number of flies crossing the line in 10 s was counted (3 trials per tube).

#### Cell immunofluorescence

Cells were seeded on optical-bottom Ibidi plates (Ibidi, Gräfelfing, Germany) and subjected to the indicated treatments. Cells were washed twice with DPBS (ThermoFisher Scientific, Waltham, MA) and fixed with 4 % paraformaldehyde (PFA) (Electron Microscopy Sciences, Morgantown, PA) for 15 min at RT, followed by three additional DBPS washes. Cells were blocked and permeabilized in DPBS (ThermoFisher Scientific, Waltham, MA) containing 5 % BSA (PAN Biotech UK Ltd, Wimborne, UK) and 0.3 % Triton X-100 (VWR, part of Avantor, Radnor, PA) for 1 h at RT. Primary antibody incubation was performed overnight at 4 °C with Arg3.1/ARC (1:500, ThermoFisher#PA5-114873, Waltham, MA) diluted in blocking buffer. After three DPBS (ThermoFisher Scientific, Waltham, MA) washes, cells were incubated with Alexa Fluor Plus-conjugated secondary antibodies (1:1000, ThermoFisher Scientific, Waltham, MA) and DAPI (ThermoFisher Scientific, Waltham, MA) for 2 h at room temperature in the dark. Following three final washes, cells were imaged using a LSM 900 confocal microscope (Carl Zeiss Microscopy GmbH, Oberkochen, Germany). Image analysis was performed using ImageJ (v2.14.0).

#### TDP-43-GFP Propagation Assay

For the TDP-43-GFP propagation experiments, U2OS cells were seeded and treated as described above. Following NaAsO_2_ (Sigma-Aldrich, St. Louis, MO) or NaAsO_2_ plus lenacapavir (MedChemExpress, Monmouth Junction, NJ) treatment, the medium was replaced with fresh culture medium for 3 h. This conditioned medium was then applied to SH-SY5Y cells for 2 h, 1 day or 2 days. Control SH-SY5Y cells were processed in parallel without exposure to non-treated U2OS conditioned medium. After the indicated incubation periods, SH-SY5Y cells were fixed as described for immunofluorescence, stained with DAPI (ThermoFisher, Waltham, MA), and imaged using an Axio Observer 7 epifluorescence microscope (Carl Zeiss Microscopy GmbH, Oberkochen, Germany). Image analysis was performed with ImageJ (v2.14.0).

#### *Drosophila* brain and thorax immunofluorescence

Flies were anaesthetised with CO_2_-enriched air on the flypad for easier manipulation under the microscope. Fly heads and thoraces were isolated and immersed in 70 % EtOH for 1 min and placed in DPBS (ThermoFisher, Waltham, MA) for dissection. Brains were dissected with fine forceps (Fine Science Tools, Foster City, CA) and both brains and thoraces were fixed with 4 % PFA (Electron Microscopy Sciences, Morgantown, PA) for 30 min. Thoraces were then immersed in liquid nitrogen for 10 s, transferred to cold DPBS (ThermoFisher, Waltham, MA) and sectioned longitudinally using a feather double edge razor blade (Aname, Madrid, Spain) attached to a blade holder (Fine Science Tools, Foster City, CA). Then, brains and thoraces were washed three times in DPBS (ThermoFisher, Waltham, MA) with 0.5 % Triton X-100 (VWR, part of Avantor, Radnor, PA) (DPBS-T) and incubated in blocking solution (5 % BSA and 0.02 % NaN_3_ in DPBS-T) for 30 min at room temperature (RT). Subsequently, samples were incubated overnight (O/N) at 4 °C with primary antibody solution in blocking solution at 4°C. After 4 × 20 min washes with DPBS-T, samples were incubated O/N at 4 °C with secondary antibodies conjugated to Alexa fluor plus fluorophores (1:1000, ThermoFisher, Waltham, MA) in blocking solution. DAPI (1:500, ThermoFisher, Waltham, MA) was also added for brains and Alexa Fluor™ 647 phalloidin (1:1000, ThermoFisher#A22287, Waltham, MA) for thoraces. Samples were mounted with ProLong™ Diamond Antifade Mountant (ThermoFisher, Waltham, MA). Images were obtained in an LS900 confocal microscope (Carl Zeiss Microscopy GmbH, Oberkochen, Germany) by z-stack scanning of whole-mount brains.

#### Lysotracker of *Drosophila* brains

For the lysotracker assay, 2-3 brains were dissected per round. An 8-well chamber slide (Ibidi, Gräfelfing, Germany) was prepared by placing a drop of DPBS (ThermoFisher, Waltham, MA) in the center of one well. The dissected brains were transferred to the well with fine forceps (Fine Science Tools, Foster City, CA). A LysoTracker staining solution was prepared in two steps: first, 1 µL of LysoTracker™ Red DND-99 (ThermoFisher, Waltham, MA) was diluted in 99 µL of DPBS (ThermoFisher, Waltham, MA) to make dilution 1; then, 5 µL of dilution 1 was mixed with 495 µL of DPBS in an opaque Eppendorf tube to make dilution 2. The DPBS (ThermoFisher, Waltham, MA) in the chamber well was carefully removed using a 200 µL LoBind pipette tip (Sigma-Aldrich, St. Louis, MO) and 200 µL of Dilution 2 was added, ensuring that the brains were detached and floating in the solution. The chamber slide was covered with aluminum foil and incubated on a shaker at 50 rpm for 7 min. After incubation, the Lysotracker solution was removed, and the brains were washed by adding 200 µL of DPBS (ThermoFisher, Waltham, MA) to the well and immediately removing it. A second wash was performed by adding a further 200 µL of DPBS (ThermoFisher, Waltham, MA) to the well, followed by incubation for 5 min on a shaker at 50 rpm, covered with aluminum foil. After the washes, the DPBS (ThermoFisher, Waltham, MA) was removed and one drop of ProLong™ Diamond Antifade Mountant (ThermoFisher, Waltham, MA) was added to the well. Images were captured in an LS900 confocal microscope (Carl Zeiss Microscopy GmbH, Oberkochen, Germany) within a maximum of 5 min due to the sensitivity of the staining and the rapid decay of fluorescence.

#### Transmission Electron Microscopy sample preparation

For TEM visualization, 4–5 brains were dissected using fine forceps (Fine Science Tools, Foster City, CA) from *Drosophila melanogaster* of the genotypes Control and Glia hTDP-43 at three time points: 1 day, 1 week, and 2 weeks. The dissected brains were immediately immersed in approximately 300 µL of fixative solution. The fixative consisted of 2.5 % glutaraldehyde and 2 % paraformaldehyde in 0.1 M phosphate buffer at pH 7.4 (Electron Microscopy Sciences, Morgantown, PA). The fixative was maintained at RT to match the sample temperature at the time of immersion. Samples were fixed at RT for 2 h with gentle agitation and then transferred to 4 °C for the remainder of the 48-hour fixation process. After fixation, samples were washed thoroughly with 0.1 M phosphate buffer. Four washes of 30 min each were performed to ensure removal of residual fixative.

The fixed brains were then sent to the Electron Microscopy Core Facility at the Principe Felipe Research Center (Valencia, Spain). Ultrathin sections were prepared using osmium tetroxide, uranyl acetate and lead citrate as contrast agents. Visualization of the prepared samples was carried out at the Electron Microscopy Department of CIC Biomagune (Donostia-San Sebastián, Spain).

#### RNA extraction, qPCR and PCR assay

Total RNA was extracted from 3-4 independent experimental pools using the RNeasy Mini Kit (QIAGEN, Hilden, Germany). Each *Drosophila* pool consisted of ≈20 brains.

Reverse transcription was performed with the High-Capacity cDNA Reverse Transcription Kit with RNase Inhibitor (ThermoFisher, Waltham, MA), following the manufacturer’s instructions.

Quantitative real-time PCR was performed on a CFX384 Touch Real-Time PCR Detection System (Bio-Rad, Hercules, CA) using Power SYBR Green Master Mix (ThermoFisher, Waltham, MA), 300 nM of primer pair, and 10 ng of cDNA. *Tubulin* was used as a housekeeping gene in *Drosophila* and *RPS17* in cells. The 2-ΔΔCt metric was used for relative quantification. The primer used to quantify *TARDBP* expression was a pre-designed sequence obtained from Sigma-Aldrich (H_TARDBP_1). For *Arc1* expression, the following primers were used: forward 5’-CATCATCGAGCACAACAACC-3’ and reverse 5’-CTACTCCTCGTGCTGCTCCT-3’, as described in Mosher *et al.* ^47^. For *ARC/Arg3.1*, the primers were: forward 5’-CCTCAGCTCCAGTGATTC-3’ and reverse 5’-TTGAGACCTGTTGTCACTC-3’. For detection of the *Dyb* cryptic exon, we used the following primers: forward 5’-CGCGATAAGTGCAGAGAAACAG-3’ and reverse 5’-

GCTATGGATGGGAATTCCGCAT-3’, previously reported in Donde *et al* ^24^ All primers were ordered from Sigma-Aldrich (St. Louis, MO).

To confirm the presence of the cryptic exon in *Dyb*, PCR was performed using the ImmoMix^TM^ reaction mix (Meridian Bioscience, Cincinnati, OH) with the previously mentioned primers. The samples were loaded onto a 1 % agarose gel and the signal was visualized using an iBright FL1000 Imaging System (ThermoFisher, Waltham, MA). The band was expected at 214 bp, as described in Donde *et. al* ^24^

#### *Drosophila* brains single-cell sample preparation

For the single cell experiment approximately 30 brains per sample were dissected with fine forceps (Fine Science Tools, Foster City, CA) in Schneider’s *Drosophila* Medium (ThermoFisher, Waltham, MA) supplemented with 0.1 µM tetrodotoxin (TTX; Bio-Techne, Minneapolis, MN), 20 µM 6,7-Dinitroquinoxaline-2,3-dione (DNQX; Santa Cruz Biotechnology, Dallas, TX) and 50 µM D(−)-2-Amino-5-phosphonovaleric acid (D-AP5; Santa Cruz Biotechnology, Dallas, TX) to avoid transcriptomic changes. Dissected brains were placed in 1.5 mL LoBind Eppendorf tubes (ThermoFisher, Waltham, MA) and washed with Schneider’s *Drosophila* Medium (ThermoFisher, Waltham, MA). Brains were incubated in 450 µL Schneider’s *Drosophila* Medium (ThermoFisher, Waltham, MA) containing papain and collagenase 1.11 mg/mL (Sigma-Aldrich, St. Louis, MO) for 30 min at RT with gentle shaking. After incubation, the brains were washed with Schneider’s *Drosophila* Medium (ThermoFisher, Waltham, MA) and mechanically dissociated in 150 µL of DPBS (ThermoFisher, Waltham, MA) + 0.01 % BSA (PAN Biotech UK Ltd, Wimborne, UK) using a 200 µL pipette with LoBind tips. To facilitate dissociation, 200 µL pipette LoBind tips were pre-rounded by heating over a flame with a 0.3 mm pin (Fine Science Tools, Foster City, CA). Separately, a 10 µm filter (PluriSelect, Leipzig, Germany) was prepared by precoating with DPBS (ThermoFisher, Waltham, MA) + 0.01 % BSA (PAN Biotech UK Ltd, Wimborne, UK) and placed over a 50 mL Falcon tube on ice. Tissue dissociation was performed in four rounds of three gentle up and down pipettings using a LoBind pipette tip set to a volume of 150 µL DPBS (ThermoFisher, Waltham, MA) containing 0.01 % BSA (PAN Biotech UK Ltd, Wimborne, UK). Care was taken to avoid the formation of bubbles. After each round, the suspension was centrifuged at 0.3 rpm for 15 s at 4 °C. The supernatant was collected after each centrifugation step and passed through the 10 µm filter. The filtered suspension was collected in a 1.5 mL LoBind Eppendorf tube.

The single-cell suspension was processed using the BD Rhapsody™ Single-Cell Analysis System (BD Biosciences, San Jose, CA). Cell concentration and viability were assessed on this system prior to pooling. The desired number of cells was adjusted to a final volume of 620 µL and the sample was prepared for further processing according to the BD Rhapsody™ protocol. (https://scomix.bd.com/hc/en-us/articles/360023044532-BD-Rhapsody-WTA-Protocols)

#### Single-cell RNAseq of *Drosophila* brains

##### Library preparation and sequencing

Single-cell cDNA libraries were generated following the manufacturer’s protocol for the BD Rhapsody Single-Cell Analysis System. Libraries were sequenced on a NovaSeq 6000 (Illumina, San Diego, CA) at Biobizkaia Health Research Institute (Spain).

##### FASTQ file preprocessing

FASTQ files were preprocessed using the BD Rhapsody™ Sequence Analysis Pipeline (detailed usage instructions are available on the <u>BD website</u>). Briefly, preprocessing was performed with *cwl-runner* v2.0 and included the following steps (LINK - https://bd-rhapsody-bioinfo-docs.genomics.bd.com/steps/steps_quality.html). (1) read quality filtering: R1/R2 pairs with low complexity (single nucleotide frequency ≥ 0.55 for R1 and ≥ 0.80 for R2) or short reads (< 60 bp for R1 and < 40 bp for R2) were removed); (2) template switch oligo removal; (3) cell label sequence (CLS) and unique molecular identifier (UMI) extraction; (4) alignment with STAR (v2.52b)^49^; (5) collapse of reads with identical CLS and UMI, as well as reads with 1-2 mismatches after recursive substitution error correction (RSEC) correction; and (6) selection of cells by applying the second derivative criterion to the distribution of sorted and log₁₀-transformed number of reads per cell.

The *Drosophila melanogaster* reference genome (version dm6; Ensembl release 111 FASTA and GTF annotations) was utilized for downstream alignment. To prevent string-parsing conflicts within the pipeline, GTF entries containing "nan" text patterns (e.g., the nanchung gene annotation) were systematically masked. The reference index was generated using the BD Rhapsody pipeline command *cwl-runner make_rhap_reference_2.0.cwl* with the modified FASTA and GTF files. Crucially, to accurately quantify reads originating from the human transgene, which consists of the human *TARDBP* coding sequence (CDS) cDNA rather than the full genomic locus, the full-length *TARDBP* CDS sequence was retrieved from Ensembl, appended to the *Drosophila* reference FASTA as an exogenous contig, and its corresponding coordinate boundaries were explicitly defined within the custom GTF annotation file.

##### Sample processing

Following the preprocessing of raw FASTQ files, h5ad AnnData files were processed in two distinct contexts: (1) a full processing pipeline applied to entire samples and (2) a partial processing pipeline tailored for specific dataset fractions or populations. For the full pipeline, Control/Glia-hTDP43 samples were processed separately from FUS samples, but the processing steps remained identical across groups.

The full pipeline consists of a first part where all samples are processed jointly using *scanpy* (v1.10.4)^50^: (1) filtering of cells expressing <200 genes and detected in <15 cells, (2) further filtering based on the distribution of reads per cell retaining cells in the [5, 95] percentile distribution, and with mitochondrial read percentages in the [0.1 - 20] range; (3) doublet removal with scrublet (v0.2.3)^51^ using an expected doublet rate of 0.05; (4) normalization across cells using *scranpy* (v0.1); and (5) log1p transformation.

Subsequently, each sample was processed individually, following these steps: (6) Principal Component Analysis (PCA) to the topmost 40 components; (7) k-Nearest

Neighbor (kNN) graph computation with a k value of 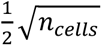 and cosine similarity; (8) highly variable gene (HVG) selection using *triku* (v2.1.6)^54^ with standard parameters; (9) refinement of PCA a and kNN graph with HVGs; (10) Uniform Manifold Approximation and Projection (UMAP, v0.5.7)^55^ for plots and (11) unsupervised clustering with *leiden* (v0.10.2)^56^.

In cases where an analysis is performed with several samples at once, *harmony* (v0.1.5)^57^ is used to correct for batch effects. *Harmony* is run after PCA and before kNN graph building.

##### Cell type assignment

Cell clusters were assigned to cell types using the *cell_assign* algorithm (v0.1.2)^59^. This algorithm maps dataset clusters to predefined populations based on a dictionary of populations and their respective markers (**Table S3**), derived from the literature (e.g. Allen *et. al*^61^, monoaminergic populations from Croset *et. al*^62^, Kenyon cell markers from Corrales *et. al*^63^) as well as by mapping DEGs to known fly atlas^61,64^. Successful mapping requires DEGs mapping to a specific population in the atlas, and DEGs from the atlas population mapping back to the original population from the dataset.

Following cell type assignment, it was observed that certain cell types, specifically Glia and Non-functional populations, required further refinement in their characterization. To address this, subsets of cells were isolated and re-processed. These subsets included glial cell types as well as basal cholinergic, glutamatergic, and GABAergic populations. Details of the specific analyses conducted for each subtype are provided in the notebooks in the ***Data availability*** section.

##### Pseudotime analysis

To perform the pseudotime analysis, specific cell subtypes were isolated and re-processed. Three tools were employed independently, each serving a distinct purpose in the analysis.

Potential of Heat-diffusion for Affinity-based Trajectory Embedding (PHATE) (v1.0.11)^66^ was first applied for dimensionality reduction to uncover putative temporal dynamics within the cell subsets. PHATE prioritizes preserving global distances over local ones, unlike methods such as UMAP. This allows for a broader view of intercellular dynamics, including potential temporal or pseudotemporal changes. For this step, the *scanpy* implementation of PHATE was used, with the number of neighbors (*k*) adjusted based on the dataset (default: 20) and the kernel tail decay (*a*) parameter set to 10.

To analyze relationships between clusters, the partition-based graph abstraction (PAGA) algorithm (v1.2)^67^ was used. PAGA constructs a graph representing relationships between predefined clusters or partitions, which can also be interpreted as a pseudotemporal trajectory.

For glial subtypes, visual inspection of the PHATE representation guided a manual annotation of clusters. This annotation incorporated information on the sample condition and collection day to refine the cluster assignments further.

To determine pseudotemporal pathways, Slingshot^68^ was applied via its Python wrapper *pyslingshot* (v0.1.5). Slingshot uses low-dimensional representations to construct branched trajectories that represent potential developmental pathways.

To ensure that pseudotemporal analyses produced results with biological relevance, all three methods—PHATE, PAGA, and Slingshot—were used in combination. Consistency among the outputs of these tools was assessed to increase confidence in the biological relevance of the trajectories.

To identify genes associated with the pseudotemporal trajectory, pseudotime values from Slingshot were used to adjust individual gene expression data with a Generalized Additive Model (GAM) via *pyGAM* (v0.9.1). A Gamma-type GAM with six splines was employed, and genes with a p-value < 10⁻⁵ were retained. For illustrative purposes, these genes were filtered further based on four criteria: minimum number of counts, maximum standard deviation, minimum mean, and maximum mean. The retained genes were ranked by their mean pseudotime position, calculated as the average pseudotime value corresponding to their expression.

#### Computational modeling of Arc1/ARC–ligand interactions

Computational analyses were performed to evaluate the ability of Lenacapavir and Bersacapavir to bind the capsid-forming interfaces of *Drosophila* Arc1 and human ARC. The crystallographic *Drosophila* Arc1 capsid assembly was obtained from the Protein Data Bank (PDB ID: 6TAS). Because no experimentally resolved hexameric structure is available for human ARC, a capsid-forming human ARC assembly was generated using AlphaFold3 and subsequently assessed by molecular dynamics simulations. Three independent 200-ns simulations were performed for both Arc1 and ARC hexamers using GROMACS 2024.4 with the AMBER ff14SB force field in explicit TIP3P water containing 150 mM KCl. For human ARC, the last 50 ns of each trajectory were combined and clustered, and the centroid of the dominant cluster was selected as the representative structure for downstream analyses.

Protein and ligand structures were prepared using Maestro/Schrödinger Release 2024 -4. Lenacapavir and Bersacapavir were processed with LigPrep, including enumeration of relevant protonation/tautomeric states while preserving stereochemistry. Minimal dimeric units containing the inter-subunit capsid-forming interface were extracted from the Arc1 and ARC assemblies and used for ligand-binding calculations. Candidate binding poses were generated by induced-fit docking at the predicted interface pocket and clustered to remove redundant conformations. Representative poses were then evaluated by binding-pose metadynamics to assess pose stability in explicit solvent and to filter out unstable binding modes.

Selected poses were further refined by 100-ns molecular dynamics simulations in Desmond, used exclusively as a stability filter, and protein–ligand interaction persistence was analyzed using the Simulation Interaction Diagram workflow. Binding modes dominated by transient or water-mediated contacts, or showing unstable ligand retention within the pocket, were excluded from further analysis. Poses that passed both the binding-pose metadynamics and Desmond filtering steps were then subjected to an independent 100-ns molecular dynamics simulation in GROMACS 2024.4, using the AMBER ff14SB force field in explicit TIP3P water. Protein and ligand RMSD and RMSF, together with protein–ligand interaction fingerprints, were monitored throughout this simulation as an additional stability check, and the final frame was used as the starting conformation for absolute binding free-energy calculations. These were performed using FEP+ with three independent replicas per selected pose. Binding free energies were converted into estimated dissociation constants to compare the relative binding of Lenacapavir and Bersacapavir to Arc1 and human ARC.

### Statistical analysis

All statistical analyses were performed using GraphPad Prism 8.0.1 software. Comparisons between two groups were conducted using two-tailed Student’s t-test. For experiments with more than two groups, one-way ANOVA was applied, while two-way ANOVA was used for experiments with two independent factors, and Mantel-Cox analysis for *Drosophila* longevity assays. All experiments were performed in biological replicates and data are presented as mean ±SEM from independent experiments (n indicated in each experiment). Significance in all figures is indicated as: *p<0.05, **p<0.01, ***p<0.001 and ****p<0.0001.

## Supporting information

All Supp figures and Legends

## RESOURCE AVAILABILITY

### Lead contact

Further information and requests for resources, reagents and fly stocks are available from the lead contact, Dr. Gorka Gerenu or Dr. Francisco Javier Gil Bea

### Materials availability

All unique/stable fly stocks and reagents generated in this study are available from the lead contact upon request.

### Data availability

- Raw count files (h5ad format) of sc-RNAseq from Drosophila brains are available at BioStudies (S-BSST2693), and FASTQ/BAM files at ENA (PRJEB107781). All intermediate files, outputs of notebooks and other files derived from this analysis are available at Zenodo (10.5281/zenodo.18496535). The raw code is available at GitHub (https://github.com/NanoNeuro/ALS_fly_scRNAseq). An interactive version of the dataset will be made available at CellxGene.
- RNA-seq data of U2OS cells following the isolated overexpression of ALS/FTD-linked factors or exposure BMAA from figure S5 are available at the GEO repository with accession number GSE327905 (https://github.com/KPlab-coding/Saez-Mas-2026-oRPs-ALS) (10.64898/2026.05.18.725994).
- All input files and execution scripts required to reproduce the molecular dynamics, docking, and free energy calculations described in this study are available at GitHub (https://github.com/BilbaoComputationalBiophysics/ARC_MD_Pipeline).

## ACKNOWLEDGEMENTS

This research was supported by the Biogipuzkoa Health Research Institute (Biogipuzkoa HRI), Ikerbasque (Basque Foundation for Science), Eusko Jaurlaritza- Basque Government and CIBER-Consorcio Centro de Investigación Biomédica en Red-(CB06/05/1126, Group 609, CiberNed), Instituto de Salud Carlos III and the Spanish Ministry of Science, Innovation and Universities / Agencia Estatal de Investigación (AEI)—European Regional Development Fund.

This work was supported by the Spanish Ministry of Science, Innovation and Universities / Agencia Estatal de Investigación (AEI)[10.13039/501100011033], under project PID2023-153029OA-I00, by “ERDF A way of making Europe”, by “ERDF/EU”, by the “European Union” or by the “European Union NextGenerationEU/PRTR”. Instituto Salud Carlos III (ISCIII) and co-funded by the European Union by ISCIII Programa Fortalece del Ministerio de Ciencia e Innovación (FORT23/00026); SEED-ALS project from ISCIII (https://www.isciii.es/en/w/estudio-seed-als-esclerosis-lateral-amiotr%C3%B3fica), by CIBERNED (CIBER de Enfermedades Neurodegenerativas, project PI19/01637, PI21/00153, PI24/1025); by Fundació La Marató (FMARATO20/001), by the Department of Education of the Basque Country through the IKUR initiative (NEURODEGENPROT); by the Department of Industry of the Basque Country through Medtech programme (MTVD22/BD/014, MTVD24/BG/006, MTVD25/BG/008) and HAZITEK programme (ZL-2025/00631, ZL-2026/00308), by Department of Health of Basque Country (projects, 2018111042, 2019222020, 2021333050, 2025333047 and 2025333026), by the Diputación Foral de Gipuzkoa (project 2021-CIEN-000020-01); by EiTB Maratoia (BIO17/ND/023/BD). AJZ, AMA, HHE, JZE, were supported by the Department of Education of the Basque Country (PRE_2020_1_0191, PRE_2022_1_0039, PRE2022_1_0315); AMA is supported by the IKUR-Nanoneuro initiative (Basque Government) and the postdoctoral fellowship from the Basque Government (POST_2025_1_0016). ASM was supported by the Department of Industry of the Basque Country BIKAINTEK Programme (011-B2/2021); LB by the Spanish National Plan for Scientific and Technical Research and Innovation -Ramon y Cajal- (RYC2018-024397-I) and IKERBASQUE (RF/2019/001) research programs; FGB by Roche Stop Fuga de cerebros(BIO19/ ROCHE/017/BD) and IKERBASQUE (PP/2022/003) research programs; GGL by Juan de la Cierva-Incorporaci.n(ISCIII, IJC2019-039965-I) and IKERBASQUE (RF/2023/010) research programs.

The authors acknowledge the technical and human support provided by SGIker (UPV/EHU, ERDF, EU), by Microscopy core facility of CICBiogune with special thank to Marta Gallego (Microscopy technician) and to Mario Soriano from the Electron Microscopy Core Facility at the Príncipe Felipe Research Center in Valencia, Spain.

Finally, the authors would like to extend their special thanks to the DalecandELA and ConELA foundations of ALS patients for their support and dedicate this work to all ALS patients and their families. The APC was funded by the same funding sources, in accordance with the authors’ affiliation with the University of the Basque Country (UPV/EHU).

## AUTHOR CONTRIBUTIONS

FGB and GGL conceived the project. AJZ performed and analyzed Drosophila experiments for Glia-hTDP-43 characterization and AJZ and SM performed hTDP-43 propagation IF determinations. AJZ, GGL and JZE optimized the sample preparation for scRNAseq. AMA performed computational analysis of scRNAseq datasets and participated in the interpretation of those results together with AJZ, FGB and GGL. AMA also wrote the parts of the manuscript involving scRNAseq analysis and interpretation. AJZ, GGL and FGB performed TEM experiments. AJZ, ASM and HHE performed double transgenic fly experiments. ASM and GGL designed and performed hTDP-43GFP *in vitro* propagation assay. LB has designed the ARC ASO for human *in vitro* experiments and participated in the edition of the manuscript. ASM and LRG performed ARC ASO treatment in *in vitro* propagation assay. MAD performed qPCR experiments to determine ARC levels in C9ORF72, FUS and TDP-43 Drosophila models. ASM, OFC and VL generated U2OS (TDP-43, C9ORF72 and hRNAP2) cell lines and performed transcriptomic analysis. MGI, MFL, RR, AL and AB performed the docking, molecular dynamics and computational calculations for Arc1, ARC structures and for Lenacapavir Binding and actively participated in the writing process of the manuscript. ASM, IJS and GGL performed different experiments to evaluate different CAMs (Lenacapavir and Bersacapavir) and Triumeq treatment experiments. AJZ, IJS, ASM and ALM, FGB, GGL wrote the manuscript and all authors discussed the results and edited the manuscript. ALM, FGB and GGL obtained funding for the project.

## SUPPLEMENTARY FIGURES

**Figure S1. Glial expression of hTDP-43 induces metabolic and synaptic alterations, related to Figure 1**

**(a)** Confocal micrographs of Control and Glia-hTDP43 brains (14 days) stained for anti-ubiquitin (red) and anti-Ref(2)p (orange), with quantification of mean staining intensity for both markers (n = 5). Scale bar, 40 μm.

**(b)** Confocal micrographs of Control and Glia-hTDP43 brains (1, 7, and 14 days) stained with LysoTracker™ Red DND-99 (red) and quantification of mean fluorescence intensity (n = 3). Scale bar, 20 μm.

**(c)** TEM images of Glia-hTDP43 brains at 7 and 14 days, highlighting the presence of defective lysosomal structures known as lamellar bodies.

**(d)** Confocal images of 7 days Control and Glia-hTDP43 thoraces stained with phalloidin (blue) and anti-nc-82 (white), with detailed images of anti-nc-82 staining along an axonal branch of both groups and quantification of density and area of muscle synaptic contacts (n = 9). Scale bar, 20 μm.

All representative images were acquired under identical conditions and magnification for each marker. Data are shown as mean ± SEM. Statistical significance: *p < 0.05, p < 0.01, ***p < 0.001, ****p < 0.0001 (Student’s t test).

**Figure S2. Glial *Arc1* activation is characteristically conserved across non-TDP-43 ALS/FTD genetic models, Related to Figures 1 and 3**.

**(a)** Schematic layout of the glia-specific human FUS expression strategy (Glia-hFUS) driven by the REPO-Gal4 system, and qPCR validation of *FUS* transgene expression in whole heads.

**(b)** Quantification of systemic locomotor decay via climbing assay at day 7 (left) and Kaplan-Meier survival curves (right).

**(c)** High-magnification confocal micrographs with representative orthogonal Z-stack projections (bottom and right panels) of 5-day-old Glia-hFUS brains stained with DAPI (blue), anti-TH (dopaminergic neurons, green), and anti–hFUS (red). Cross-hairs highlight the non-cell-autonomous internalization of glia-derived hFUS protein within recipient TH⁺ neuronal cell bodies. Scale bar, 5 μm.

**(d)** UMAP plot of Glia-hFUS 1 day sample, color-coded by major cell populations.

**(e)** High-resolution subclustering UMAP of the isolated glial compartment from 1-day-old Glia-hFUS brains (GlA1, GlEn, GlSu) overlaid with a continuous feature expression gradient, showing robust, early *Arc1* transcript upregulation across specific pathology-associated glial subsets.

**(f)** UMAP plots illustrating *Arc1* expression of all cell types in Glia-hFUS dataset.

**(g)** qPCR analysis of *Arc1* expression levels in Control and Glia-hFUS models at 1 day (n = 3).

**(h)** Schematic representation of the glia-targeted C9orf72 repeat expansion model (Glia-hC9orf72; 49xG_4_C_2_), alongside matching physiological characterizations showing day 7 locomotor performance (climbing assay), longitudinal lifespan survival curves, and day 7 qPCR validation of mass *Arc1* transcript upregulation in fly brains (n = 3).

All representative images were acquired under identical conditions and magnification. Data are shown as mean ± SEM. Statistical significance: *p < 0.05, p < 0.01, ***p < 0.001, ****p < 0.0001 (Student’s t test and Mantel-Cox for the longevity assay).

**Figure S3. UMAP subpopulation analysis and functional GO terms of Glia-hTDP43 datasets, related to Figure 3**.

**(a)** Longitudinal feature expression UMAP plots tracking the transcript levels of canonical neuronal identity markers: *ChAT* (cholinergic; green vertical bar), *VGlut* (glutamatergic; red vertical bar), and *Gad1* (GABAergic; purple vertical bar) across a temporal continuum (1, 7, and 14 days post-eclosion).

**(b)** Comparative scatter plot profiling GO terms derived from the transcriptional signature of the emergent non-functional neuronal cluster. The coordinates map the statistical significance (-log_10_[adjusted p-value]) of identified functional families in Control (x-axis) versus Glia-hTDP43 datasets (y-axis). The dashed diagonal line denotes structural equivalence between conditions, isolating specific macromolecular and translational inhibitory pathways that are selectively accelerated or amplified under hTDP-43 proteotoxic stress.

**Figure S4. Pseudotemporal heatmaps of gene expression across glial branches, related to Figure 4**.

Heatmaps of gene expression sorted pseudotemporally (t0 on the left, t1 on the right). Genes are sorted based on the mean ranking of cells that have a positive (greater than 0) expression. **(a)** Astrocyte-like glia branch. **(b)** Ensheathing glia branch.

**Figure S5. Volcano plot matrices identify *ARC* as a convergent, transcriptionally invariant hub across heterogeneous genetic and environmental ALS/FTD stressors, related to Figure 4**.

Volcano plots showing differential gene expression profiles from U2OS cells subject to different pathological contexts (TDP-43, PR97 poly-dipeptide repeats, and mutant hnRNPA2) alongside a kinetic environmental stress axis utilizing the cyanobacterial neurotoxin beta-methylamino-L-alanine (BMAA 6h and BMAA 24h). Across all cases, the *ARC* transcript (highlighted in red) consistently xdisplays robust, statistically significant upregulation (log_2_FC > 0).

**Figure S6. Lenacapavir binding to HIV-1 capsid. Related to Figure 5**.

**(a)** Hexameric representation of the HIV–Lenacapavir complex (PDB: 6V2F). Each monomer is colored uniquely, and Lenacapavir is shown in orange.

**(b)** Dimeric representation of the E–F interface, highlighting the monomer subunits (grey and green) and Lenacapavir at the inter-subunit interface.

**(c)** Detailed interactions of Lenacapavir within HIV after 100 ns Desmond molecular dynamic (MD) simulation. Purple arrows indicate hydrogen bonds: arrows originating from a residue represent donors, whereas arrows pointing toward a residue indicate acceptors. Red lines correspond to π–cation interactions, and gradient blue–red lines indicate salt bridges.

**(d)** Binding Pose Metadynamics (BPMD) simulation of the HIV–Lenacapavir dimer. The pose remains highly stable throughout the simulation, with PoseScore and PerScore metrics falling within the ranges defined in Methods.

**Figure S7. Structural validation and molecular dynamics simulations of the ARC hexamer, related to Figure 5**.

**(a)** AlphaFold3 prediction of the hexameric human ARC structure. Colors represent the predicted local confidence (pLDDT), indicating the expected agreement with an experimental structure. Blue indicates high confidence, yellow low confidence, and brown very low confidence.

**(b)** AlphaFold3 prediction showing only the capsid-forming region of ARC (residues 206–361).

**(c)** Root mean square deviation (RMSD) from three independent 200 ns molecular dynamics simulations of Arc1 and ARC hexamer structures performed using the AMBER ff14SB force field.

**(d)** Representative ARC hexamer structure obtained from clustering of the last 50 ns of the three independent MD trajectories.

**Figure S8. Binding pose metadynamics (BPMD) results for Arc1 and ARC complexes with Lenacapavir and Bersacapavir. Related to Figure 6**.

**(a)** Arc1–Lenacapavir. **(b)** ARC–Lenacapavir. **(c)** Arc1–Bersacapavir. **(d)** ARC–Bersacapavir. Each panel summarizes the BPMD evaluation of representative poses obtained from IFD, with 20, 26, 28, and

