## Supplementary material for "Modulation of retroviral capsid assembly halts ARC-mediated TDP-43 intercellular spreading": All Supp figures and Legends

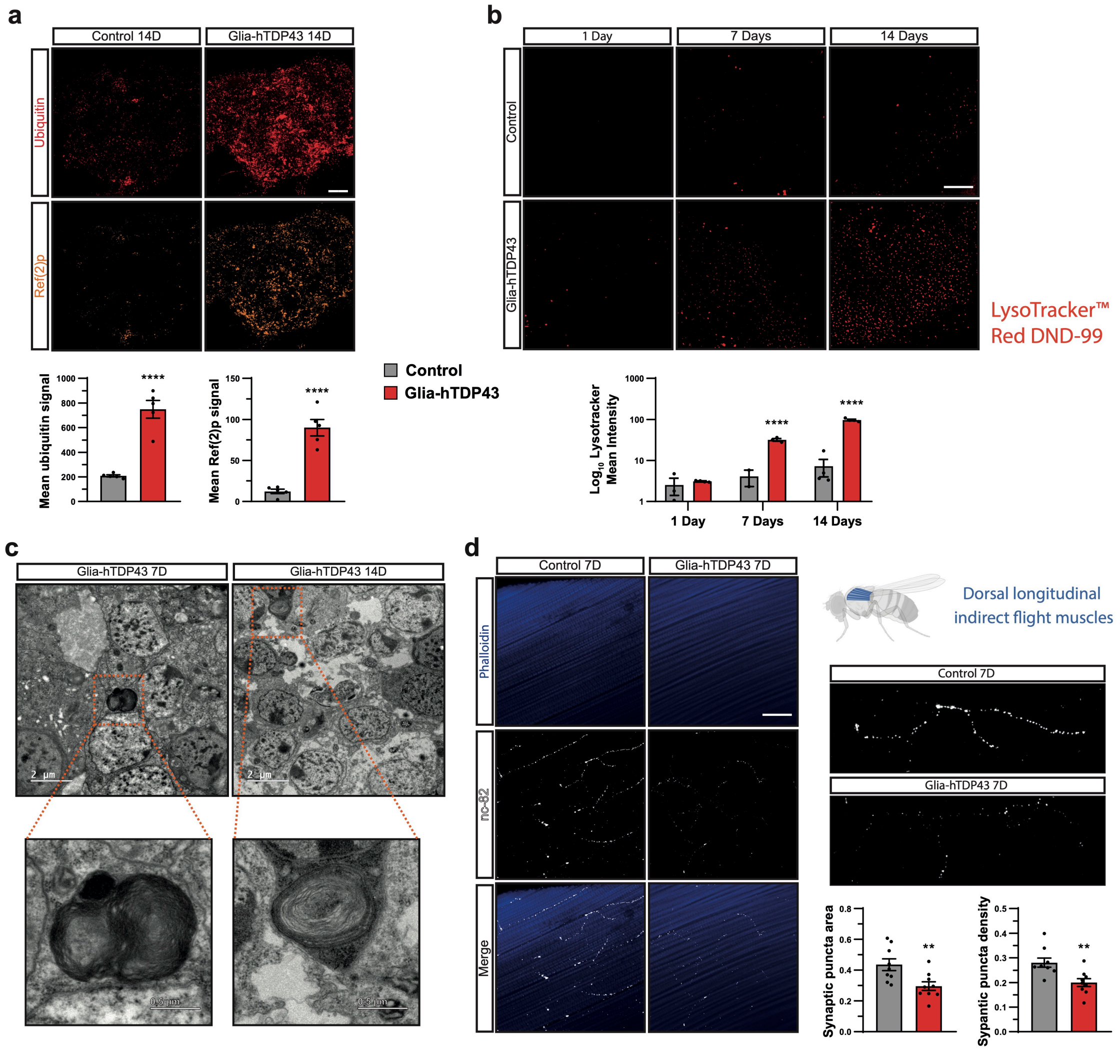
a
Control 14D
Glia-hTDP43 14D
Ubiquitin
Ref(2)p
Mean ubiquitin signal
Mean Ref(2)p signal
****
Control
Glia-hTDP43
b
1 Day
7 Days
14 Days
Control
Glia-hTDP43
LysoTracker™ Red DND-99
Log10 Lysotracker Mean Intensity
c
Glia-hTDP43 7D
Glia-hTDP43 14D
2 µm
d
Control 7D
Glia-hTDP43 7D
Phalloidin
nc-82
Merge
Dorsal longitudinal indirect flight muscles
Synaptic puncta area
Sypantic puncta density
**

**Figure S1. Glial expression of hTDP-43 induces metabolic and synaptic alterations, related to Figure 1**. **(a)** Confocal micrographs of Control and Glia-hTDP43 brains (14 days) stained for anti-ubiquitin (red) and anti-Ref(2)p (orange), with quantification of mean staining intensity for both markers (n = 5). Scale bar = 40 μm. **(b)** Confocal micrographs of Control and Glia-hTDP43 brains (1, 7, and 14 days) stained with LysoTracker™ Red DND-99 (red) and quantification of mean fluorescence intensity (n = 3). Scale bar = 20 μm. **(c)** TEM images of Glia-hTDP43 brains at 7 and 14 days, highlighting the presence of defective lysosomal structures known as lamellar bodies. **(d)** Confocal images of 7 days Control and Glia-hTDP43 thoraces stained with phalloidin (blue) and anti-nc-82 (white), with detailed images of anti-nc-82 staining along an axonal branch of both groups and quantification of density and area of muscle synaptic contacts (n = 9). Scale bar = 20 μm. All representative images were acquired under identical conditions and magnification for each marker. Data are shown as mean ± SEM. Statistical significance: *$p < 0.05$, $p < 0.01$, ***$p < 0.001$, ****$p < 0.0001$ (Student's t test).

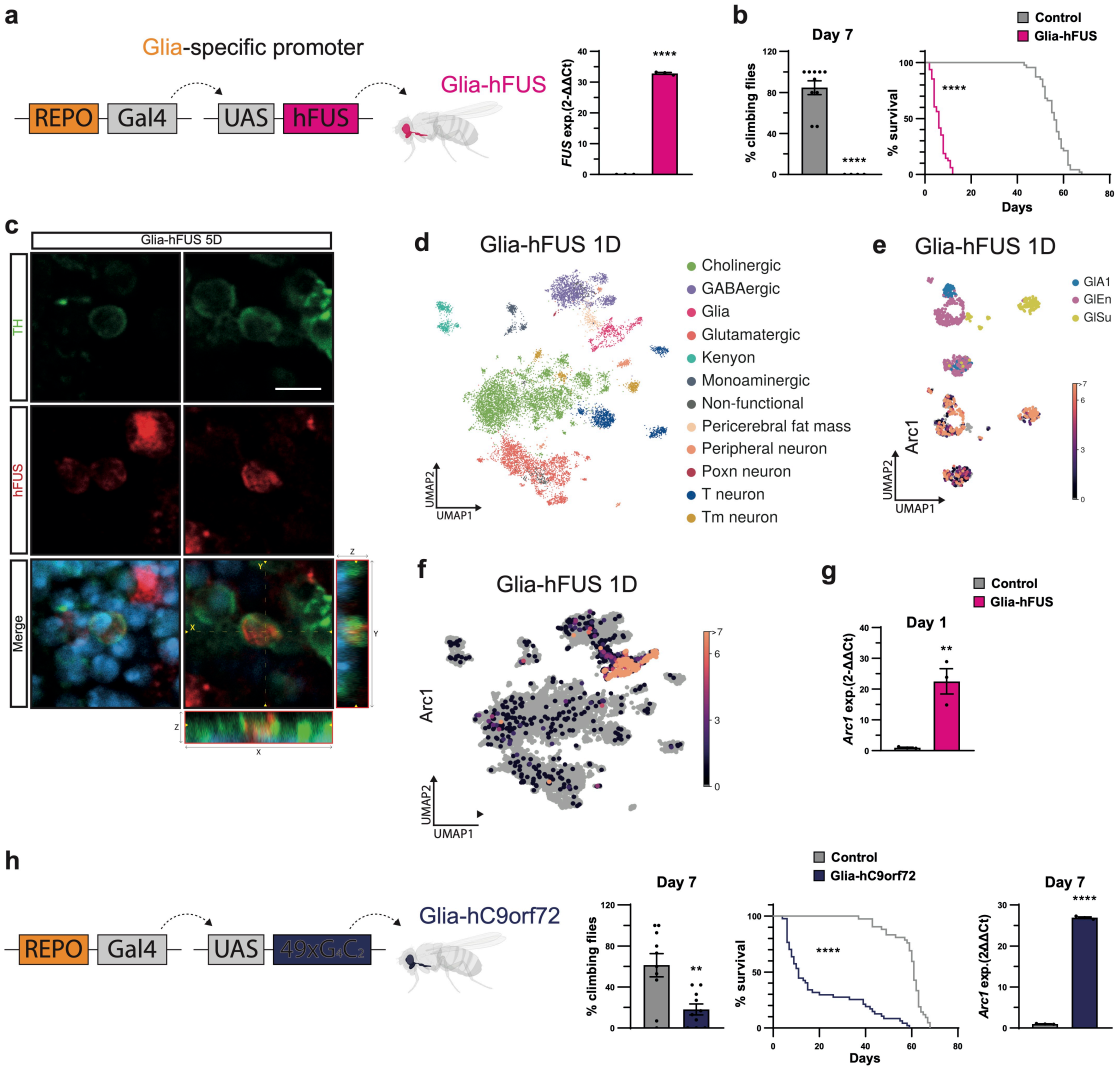

a
Glia-specific promoter
REPO
Gal4
UAS
hFUS
Glia-hFUS
FUS exp.(2-ΔΔCt)
****
b
Day 7
% climbing flies
****
Control
Glia-hFUS
% survival
****
Days
c
Glia-hFUS 5D
TH
hFUS
Merge
d
Glia-hFUS 1D
Cholinergic
GABAergic
Glia
Glutamatergic
Kenyon
Monoaminergic
Non-functional
Pericerebral fat mass
Peripheral neuron
Poxn neuron
T neuron
Tm neuron
UMAP2
UMAP1
e
Glia-hFUS 1D
GlA1
GlEn
GlSu
Arc1
UMAP2
UMAP1
f
Glia-hFUS 1D
Arc1
UMAP2
UMAP1
g
Control
Glia-hFUS
Day 1
Arc1 exp.(2-ΔΔCt)
**
h
REPO
Gal4
UAS
49xG4C2
Glia-hC9orf72
Day 7
% climbing flies
**
Control
Glia-hC9orf72
% survival
****
Days
Day 7
Arc1 exp.(2ΔΔCt)
****

**Figure S2. Glial *Arc1* activation is characteristically conserved across non-TDP-43 ALS/FTD genetic models, Related to Figures 1 and 3. (a)** Schematic layout of the glia-specific human FUS expression strategy (Glia-hFUS) driven by the REPO-Gal4 system, and qPCR validation of *FUS* transgene expression in whole heads. **(b)** Quantification of systemic locomotor decay via climbing assay at day 7 (left) and Kaplan-Meier survival curves (right). **(c)** High-magnification confocal micrographs with representative orthogonal Z-stack projections (bottom and right panels) of 5-day-old Glia-hFUS brains stained with DAPI (blue), anti-TH (dopaminergic neurons, green), and anti–hFUS (red). Cross-hairs highlight the non-cell-autonomous internalization of glia-derived hFUS protein within recipient $TH^+$ neuronal cell bodies. Scale bar, 5 µm. **(d)** UMAP plot of Glia-hFUS 1 day sample, color-coded by major cell populations. **(e)** High-resolution subclustering UMAP of the isolated glial compartment from 1-day-old Glia-hFUS brains (GlA1, GlEn, GlSu) overlaid with a continuous feature expression gradient, showing robust, early *Arc1* transcript upregulation across specific pathology-associated glial subsets. **(f)** UMAP plots illustrating *Arc1* expression of all cell types in Glia-hFUS dataset. **(g)** qPCR analysis of *Arc1* expression levels in Control and Glia-hFUS models at 1 day (n = 3). **(h)** Schematic representation of the glia-targeted C9orf72 repeat expansion model (Glia-hC9orf72; $49xG_4C_2$), alongside matching physiological characterizations showing day 7 locomotor performance (climbing assay), longitudinal lifespan survival curves, and day 7 qPCR validation of mass *Arc1* transcript upregulation in fly brains (n = 3). All representative images were acquired under identical conditions and magnification. Data are shown as mean ± SEM. Statistical significance: *p < 0.05, p < 0.01, ***p < 0.001, ****p < 0.0001 (Student’s t test and Mantel-Cox for the longevity assay).

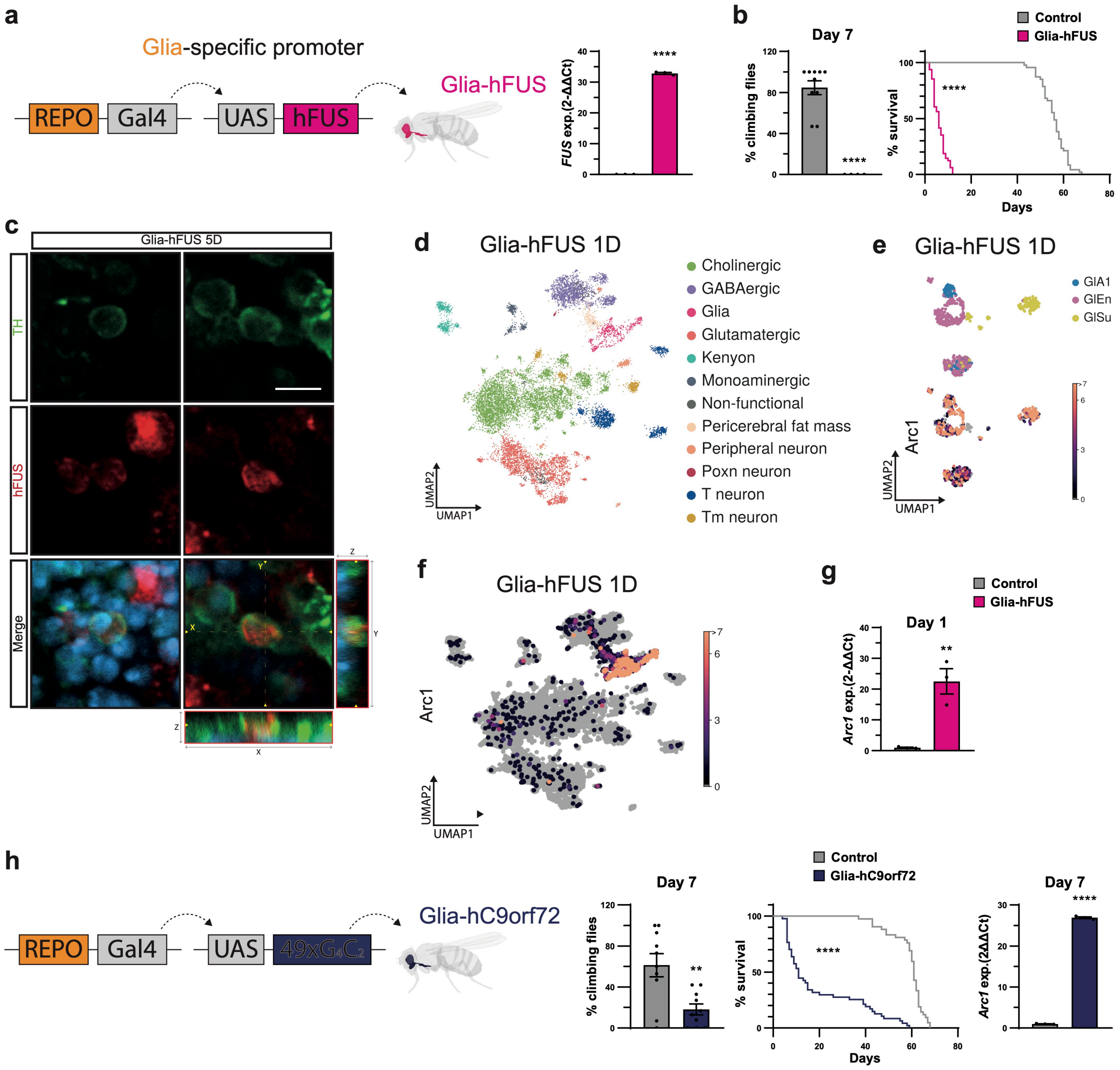

a
Glia-specific promoter
REPO
Gal4
UAS
hFUS
Glia-hFUS
FUS exp.(2-ΔΔCt)
****
b
Day 7
% climbing flies
****
Control
Glia-hFUS
% survival
****
Days
c
Glia-hFUS 5D
TH
hFUS
Merge
d
Glia-hFUS 1D
Cholinergic
GABAergic
Glia
Glutamatergic
Kenyon
Monoaminergic
Non-functional
Pericerebral fat mass
Peripheral neuron
Poxn neuron
T neuron
Tm neuron
UMAP2
UMAP1
e
Glia-hFUS 1D
GlA1
GlEn
GlSu
Arc1
UMAP2
UMAP1
f
Glia-hFUS 1D
Arc1
UMAP2
UMAP1
g
Control
Glia-hFUS
Day 1
Arc1 exp.(2-ΔΔCt)
**
h
REPO
Gal4
UAS
Glia-hC9orf72
Day 7
% climbing flies
**
Control
Glia-hC9orf72
% survival
****
Days
Day 7
Arc1 exp.(2ΔΔCt)
****

**Figure S3. UMAP subpopulation analysis and functional GO terms of Glia-hTDP43 datasets, related to Figure 3. (a)** Longitudinal feature expression UMAP plots tracking the transcript levels of canonical neuronal identity markers: *ChAT* (cholinergic; green vertical bar), *VGlut* (glutamatergic; red vertical bar), and *Gad1* (GABAergic; purple vertical bar) across a temporal continuum (1, 7, and 14 days post-eclosion). **(b)** Comparative scatter plot profiling GO terms derived from the transcriptional signature of the emergent non-functional neuronal cluster. The coordinates map the statistical significance ($-\log_{10}$[adjusted p-value]) of identified functional families in Control (x-axis) versus Glia-hTDP43 datasets (y-axis). The dashed diagonal line denotes structural equivalence between conditions, isolating specific macromolecular and translational inhibitory pathways that are selectively accelerated or amplified under hTDP-43 proteotoxic stress.

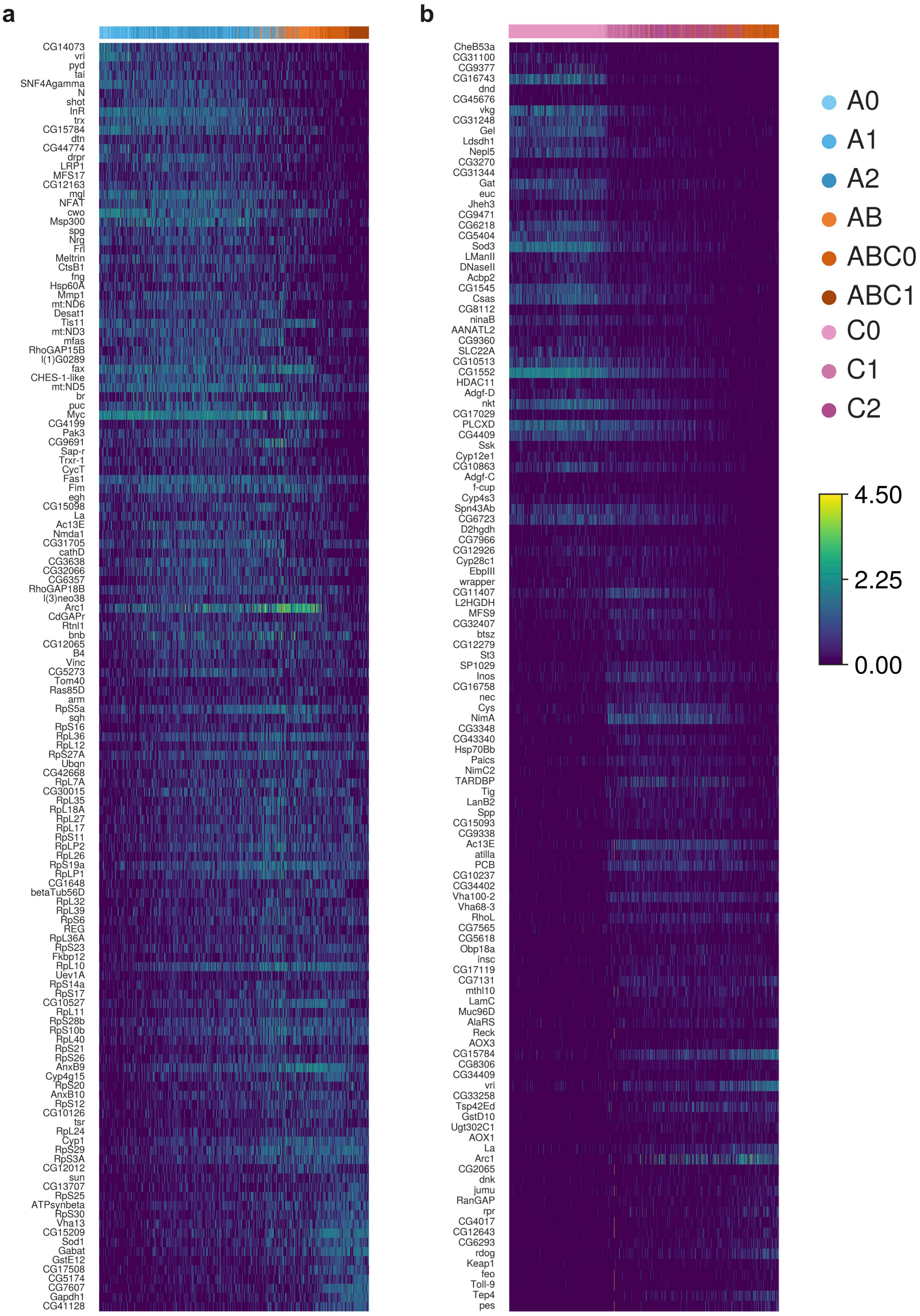
a
CG14073
vri
pyd
tai
SNF4Agamma
N
shot
InR
trx
CG15784
dtn
CG44774
drpr
LRP1
MFS17
CG12163
mgl
NFAT
cwo
Msp300
spg
Nrg
Frl
Meltrin
CtsB1
fng
Hsp60A
Mmp1
mt:ND6
Desat1
Tis11
mt:ND3
mfas
RhoGAP15B
l(1)G0289
fax
CHES-1-like
mt:ND5
br
puc
Myc
CG4199
Pak3
CG9691
Sap-r
Trxr-1
CycT
Fas1
Fim
egh
CG15098
La
Ac13E
Nmda1
CG31705
cathD
CG3638
CG32066
CG6357
RhoGAP18B
l(3)neo38
Arc1
CdGAPr
Rtnl1
bnb
CG12065
B4
Vinc
CG5273
Tom40
Ras85D
arm
RpS5a
sqh
RpS16
RpL36
RpL12
RpS27A
Ubqn
CG42668
RpL7A
CG30015
RpL35
RpL18A
RpL27
RpL17
RpS11
RpLP2
RpL26
RpS19a
RpLP1
CG1648
betaTub56D
RpL32
RpL39
RpS6
REG
RpL36A
RpS23
Fkbp12
RpL10
Uev1A
RpS14a
RpS17
CG10527
RpL11
RpS28b
RpS10b
RpL40
RpS21
RpS26
AnxB9
Cyp4g15
RpS20
AnxB10
RpS12
CG10126
tsr
RpL24
Cyp1
RpS29
RpS3A
CG12012
sun
CG13707
RpS25
ATPsynbeta
RpS30
Vha13
CG15209
Sod1
Gabat
GstE12
CG17508
CG5174
CG7607
Gapdh1
CG41128
b
CheB53a
CG31100
CG9377
CG16743
dnd
CG45676
vkg
CG31248
Gel
Ldsdh1
Nepl5
CG3270
CG31344
Gat
euc
Jheh3
CG9471
CG6218
CG5404
Sod3
LManII
DNaseII
Acbp2
CG1545
Csas
CG8112
ninaB
AANATL2
CG9360
SLC22A
CG10513
CG1552
HDAC11
Adgf-D
nkt
CG17029
PLCXD
CG4409
Ssk
Cyp12e1
CG10863
Adgf-C
f-cup
Cyp4s3
Spn43Ab
CG6723
D2hgdh
CG7966
CG12926
Cyp28c1
EbpIII
wrapper
CG11407
L2HGDH
MFS9
CG32407
btsz
CG12279
St3
SP1029
Inos
CG16758
nec
Cys
NimA
CG3348
CG43340
Hsp70Bb
Paics
NimC2
TARDBP
Tig
LanB2
Spp
CG15093
CG9338
Ac13E
atilla
PCB
CG10237
CG34402
Vha100-2
Vha68-3
RhoL
CG7565
CG5618
Obp18a
insc
CG17119
CG7131
mthl10
LamC
Muc96D
AlaRS
Reck
AOX3
CG15784
CG8306
CG34409
vri
CG33258
Tsp42Ed
GstD10
Ugt302C1
AOX1
La
Arc1
CG2065
dnk
jumu
RanGAP
rpr
CG4017
CG12643
CG6293
rdog
Keap1
feo
Toll-9
Tep4
pes
A0
A1
A2
AB
ABC0
ABC1
C0
C1
C2
4.50
2.25
0.00

**Figure S4. Pseudotemporal heatmaps of gene expression across glial branches, related to Figure 4.** Heatmaps of gene expression sorted pseudotemporally (t0 on the left, t1 on the right). Genes are sorted based on the mean ranking of cells that have a positive (greater than 0) expression. **(a)** Astrocyte-like glia branch. **(b)** Ensheathing glia branch.

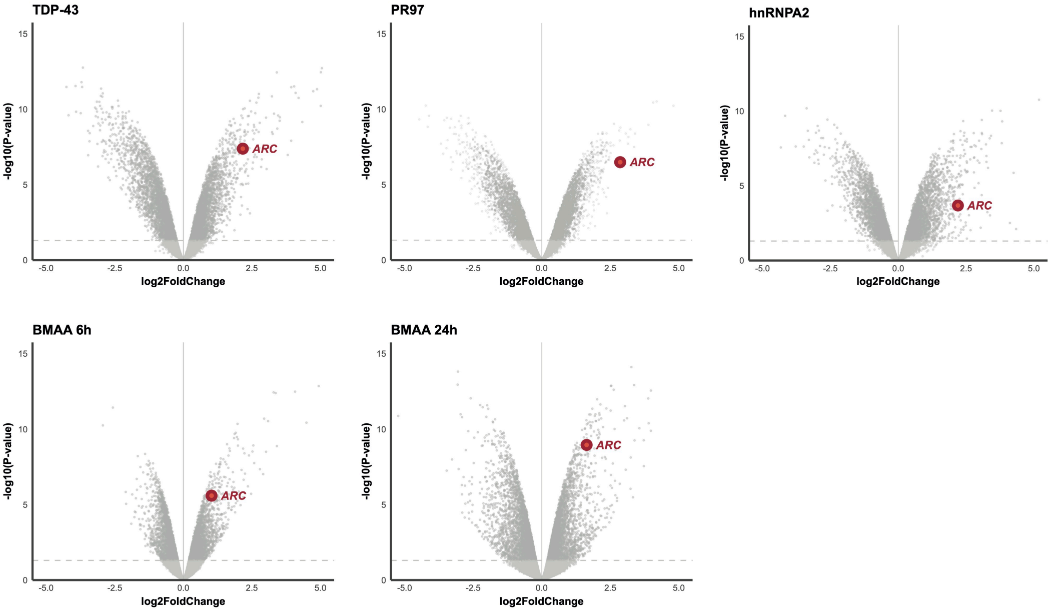

TDP-43
PR97
hnRNPA2
BMAA 6h
BMAA 24h
ARC
-log10(P-value)
log2FoldChange

**Figure S5. Volcano plot matrices identify *ARC* as a convergent, transcriptionally invariant hub across heterogeneous genetic and environmental ALS/FTD stressors, related to Figure 4.** Volcano plots showing differential gene expression profiles from U2OS cells subject to different pathological contexts (TDP-43, PR97 poly-dipeptide repeats, and mutant hnRNPA2) alongside a kinetic environmental stress axis utilizing the cyanobacterial neurotoxin beta-methylamino-L-alanine (BMAA 6h and BMAA 24h). Across all cases, the *ARC* transcript (highlighted in red) consistently displays robust, statistically significant upregulation ($log_2FC > 0$).

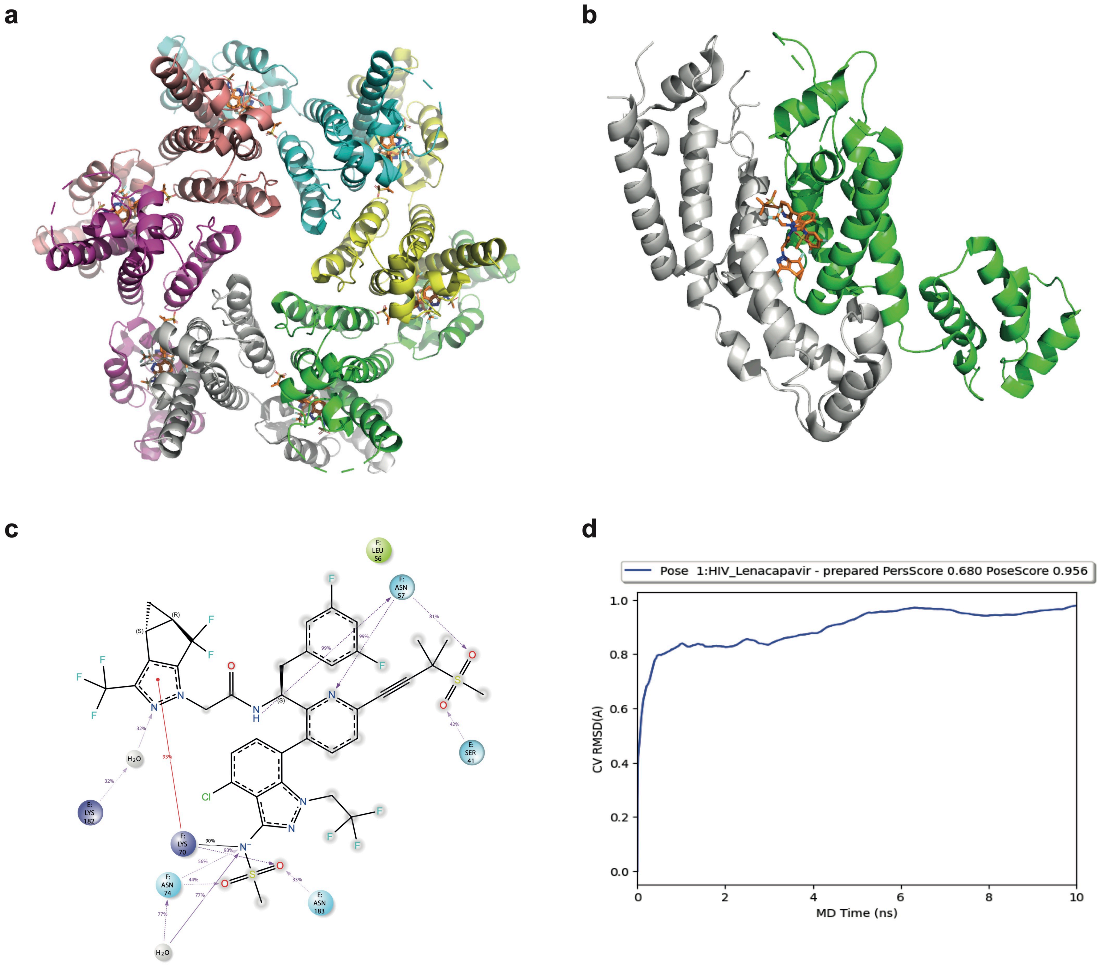
a
b
c
d
Pose 1:HIV_Lenacapavir - prepared PersScore 0.680 PoseScore 0.956
CV RMSD(A)
MD Time (ns)
0.0
0.2
0.4
0.6
0.8
1.0
0
2
4
6
8
10

**Figure S6. Lenacapavir binding to HIV-1 capsid. Related to Figure 5. (a)** Hexameric representation of the HIV–Lenacapavir complex (PDB: 6V2F). Each monomer is colored uniquely, and Lenacapavir is shown in orange. **(b)** Dimeric representation of the E–F interface, highlighting the monomer subunits (grey and green) and Lenacapavir at the inter-subunit interface. **(c)** Detailed interactions of Lenacapavir within HIV after 100 ns Desmond molecular dynamic (MD) simulation. Purple arrows indicate hydrogen bonds: arrows originating from a residue represent donors, whereas arrows pointing toward a residue indicate acceptors. Red lines correspond to π–cation interactions, and gradient blue–red lines indicate salt bridges. **(d)** Binding Pose Metadynamics (BPMD) simulation of the HIV–Lenacapavir dimer. The pose remains highly stable throughout the simulation, with PoseScore and PerScore metrics falling within the ranges defined in Methods.

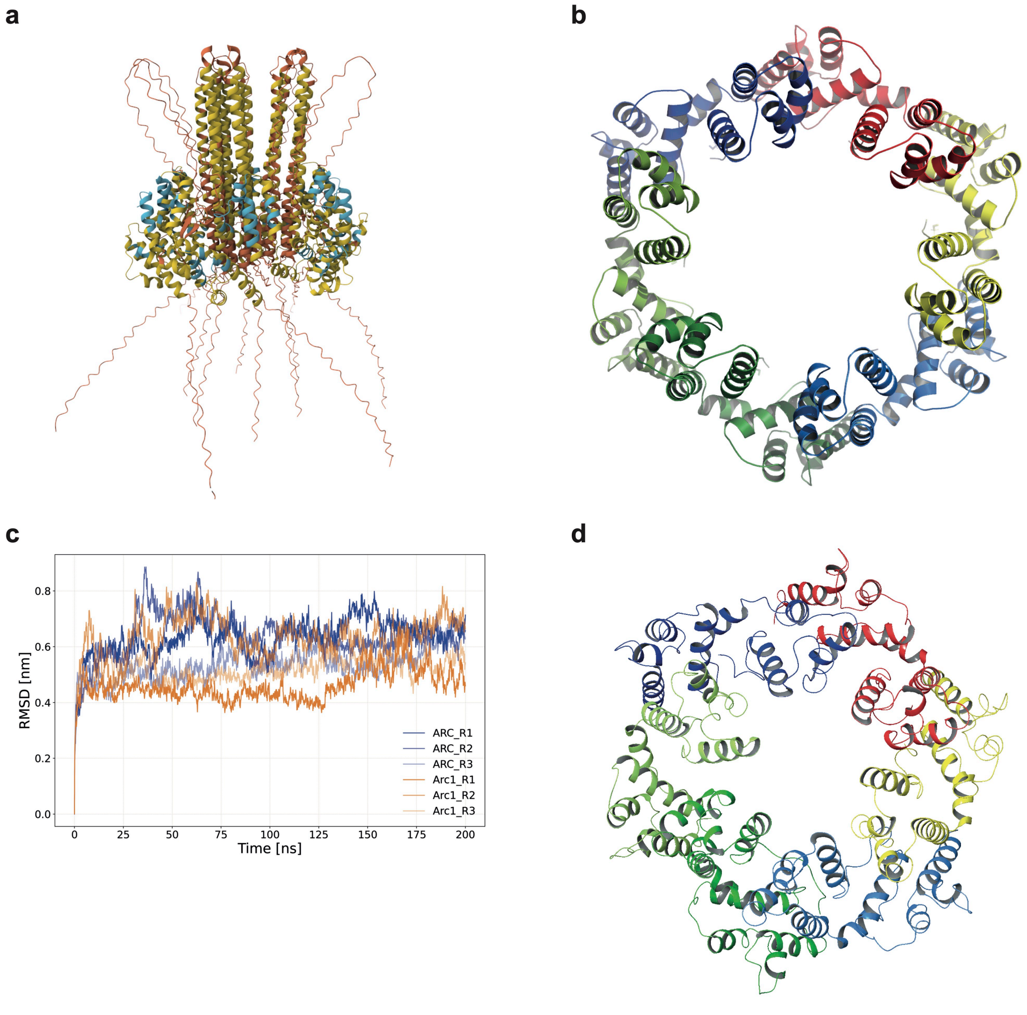
a
b
c
d
RMSD [nm]
Time [ns]
0.0
0.2
0.4
0.6
0.8
0
25
50
75
100
125
150
175
200
ARC_R1
ARC_R2
ARC_R3
Arc1_R1
Arc1_R2
Arc1_R3

**Figure S7. Structural validation and molecular dynamics simulations of the ARC hexamer, related to Figure 5. (a)** AlphaFold3 prediction of the hexameric human ARC structure. Colors represent the predicted local confidence (pLDDT), indicating the expected agreement with an experimental structure. Blue indicates high confidence, yellow low confidence, and brown very low confidence. **(b)** AlphaFold3 prediction showing only the capsid-forming region of ARC (residues 206–361). **(c)** Root mean square deviation (RMSD) from three independent 200 ns molecular dynamics simulations of Arc1 and ARC hexamer structures performed using the AMBER ff14SB force field. **(d)** Representative ARC hexamer structure obtained from clustering of the last 50 ns of the three independent MD trajectories.

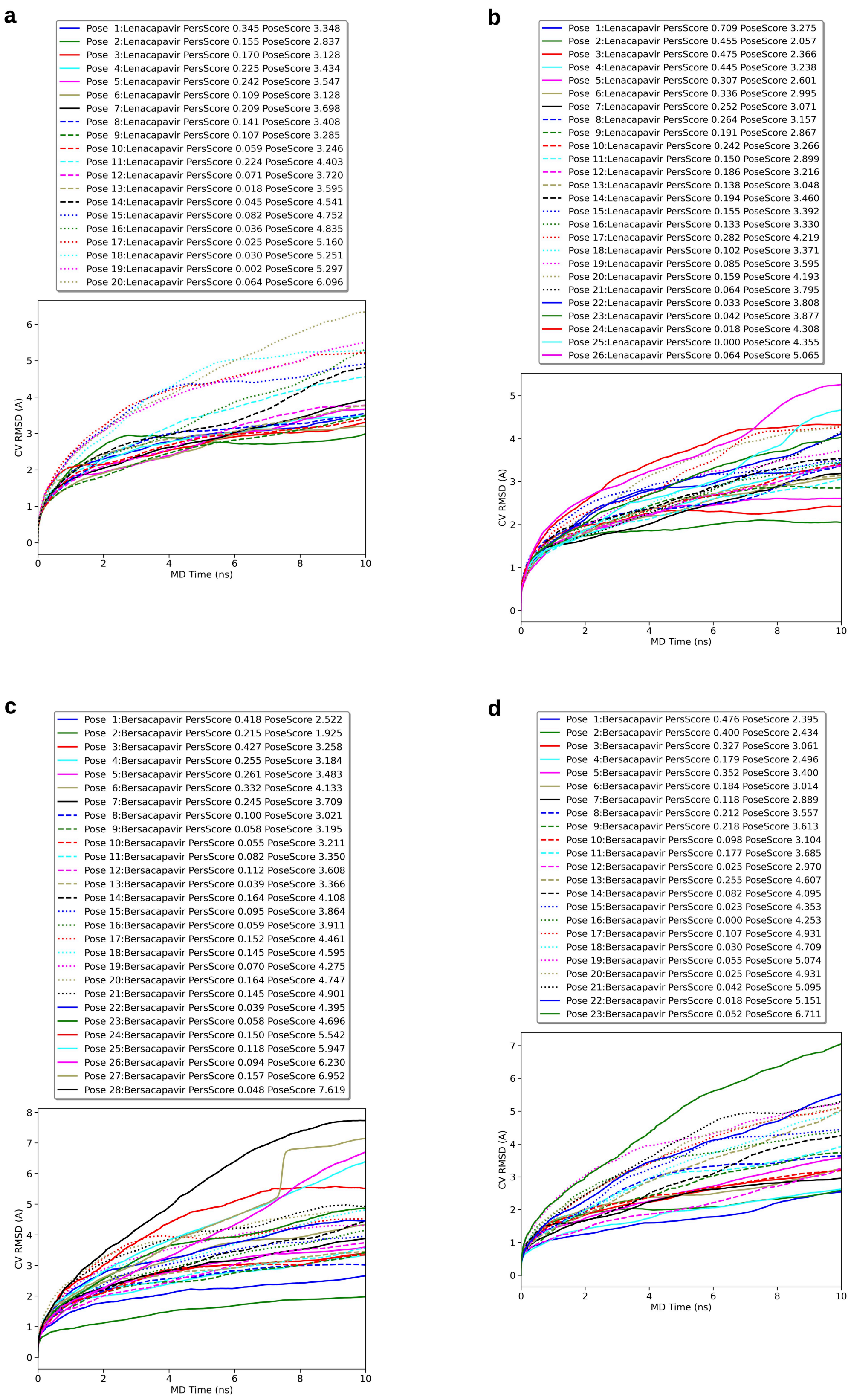
a
Pose 1:Lenacapavir PersScore 0.345 PoseScore 3.348
Pose 2:Lenacapavir PersScore 0.155 PoseScore 2.837
Pose 3:Lenacapavir PersScore 0.170 PoseScore 3.128
Pose 4:Lenacapavir PersScore 0.225 PoseScore 3.434
Pose 5:Lenacapavir PersScore 0.242 PoseScore 3.547
Pose 6:Lenacapavir PersScore 0.109 PoseScore 3.128
Pose 7:Lenacapavir PersScore 0.209 PoseScore 3.698
Pose 8:Lenacapavir PersScore 0.141 PoseScore 3.408
Pose 9:Lenacapavir PersScore 0.107 PoseScore 3.285
Pose 10:Lenacapavir PersScore 0.059 PoseScore 3.246
Pose 11:Lenacapavir PersScore 0.224 PoseScore 4.403
Pose 12:Lenacapavir PersScore 0.071 PoseScore 3.720
Pose 13:Lenacapavir PersScore 0.018 PoseScore 3.595
Pose 14:Lenacapavir PersScore 0.045 PoseScore 4.541
Pose 15:Lenacapavir PersScore 0.082 PoseScore 4.752
Pose 16:Lenacapavir PersScore 0.036 PoseScore 4.835
Pose 17:Lenacapavir PersScore 0.025 PoseScore 5.160
Pose 18:Lenacapavir PersScore 0.030 PoseScore 5.251
Pose 19:Lenacapavir PersScore 0.002 PoseScore 5.297
Pose 20:Lenacapavir PersScore 0.064 PoseScore 6.096
CV RMSD (A)
MD Time (ns)
b
Pose 1:Lenacapavir PersScore 0.709 PoseScore 3.275
Pose 2:Lenacapavir PersScore 0.455 PoseScore 2.057
Pose 3:Lenacapavir PersScore 0.475 PoseScore 2.366
Pose 4:Lenacapavir PersScore 0.445 PoseScore 3.238
Pose 5:Lenacapavir PersScore 0.307 PoseScore 2.601
Pose 6:Lenacapavir PersScore 0.336 PoseScore 2.995
Pose 7:Lenacapavir PersScore 0.252 PoseScore 3.071
Pose 8:Lenacapavir PersScore 0.264 PoseScore 3.157
Pose 9:Lenacapavir PersScore 0.191 PoseScore 2.867
Pose 10:Lenacapavir PersScore 0.242 PoseScore 3.266
Pose 11:Lenacapavir PersScore 0.150 PoseScore 2.899
Pose 12:Lenacapavir PersScore 0.186 PoseScore 3.216
Pose 13:Lenacapavir PersScore 0.138 PoseScore 3.048
Pose 14:Lenacapavir PersScore 0.194 PoseScore 3.460
Pose 15:Lenacapavir PersScore 0.155 PoseScore 3.392
Pose 16:Lenacapavir PersScore 0.133 PoseScore 3.330
Pose 17:Lenacapavir PersScore 0.282 PoseScore 4.219
Pose 18:Lenacapavir PersScore 0.102 PoseScore 3.371
Pose 19:Lenacapavir PersScore 0.085 PoseScore 3.595
Pose 20:Lenacapavir PersScore 0.159 PoseScore 4.193
Pose 21:Lenacapavir PersScore 0.064 PoseScore 3.795
Pose 22:Lenacapavir PersScore 0.033 PoseScore 3.808
Pose 23:Lenacapavir PersScore 0.042 PoseScore 3.877
Pose 24:Lenacapavir PersScore 0.018 PoseScore 4.308
Pose 25:Lenacapavir PersScore 0.000 PoseScore 4.355
Pose 26:Lenacapavir PersScore 0.064 PoseScore 5.065
CV RMSD (A)
MD Time (ns)
c
Pose 1:Bersacapavir PersScore 0.418 PoseScore 2.522
Pose 2:Bersacapavir PersScore 0.215 PoseScore 1.925
Pose 3:Bersacapavir PersScore 0.427 PoseScore 3.258
Pose 4:Bersacapavir PersScore 0.255 PoseScore 3.184
Pose 5:Bersacapavir PersScore 0.261 PoseScore 3.483
Pose 6:Bersacapavir PersScore 0.332 PoseScore 4.133
Pose 7:Bersacapavir PersScore 0.245 PoseScore 3.709
Pose 8:Bersacapavir PersScore 0.100 PoseScore 3.021
Pose 9:Bersacapavir PersScore 0.058 PoseScore 3.195
Pose 10:Bersacapavir PersScore 0.055 PoseScore 3.211
Pose 11:Bersacapavir PersScore 0.082 PoseScore 3.350
Pose 12:Bersacapavir PersScore 0.112 PoseScore 3.608
Pose 13:Bersacapavir PersScore 0.039 PoseScore 3.366
Pose 14:Bersacapavir PersScore 0.164 PoseScore 4.108
Pose 15:Bersacapavir PersScore 0.095 PoseScore 3.864
Pose 16:Bersacapavir PersScore 0.059 PoseScore 3.911
Pose 17:Bersacapavir PersScore 0.152 PoseScore 4.461
Pose 18:Bersacapavir PersScore 0.145 PoseScore 4.595
Pose 19:Bersacapavir PersScore 0.070 PoseScore 4.275
Pose 20:Bersacapavir PersScore 0.164 PoseScore 4.747
Pose 21:Bersacapavir PersScore 0.145 PoseScore 4.901
Pose 22:Bersacapavir PersScore 0.039 PoseScore 4.395
Pose 23:Bersacapavir PersScore 0.058 PoseScore 4.696
Pose 24:Bersacapavir PersScore 0.150 PoseScore 5.542
Pose 25:Bersacapavir PersScore 0.118 PoseScore 5.947
Pose 26:Bersacapavir PersScore 0.094 PoseScore 6.230
Pose 27:Bersacapavir PersScore 0.157 PoseScore 6.952
Pose 28:Bersacapavir PersScore 0.048 PoseScore 7.619
CV RMSD (A)
MD Time (ns)
d
Pose 1:Bersacapavir PersScore 0.476 PoseScore 2.395
Pose 2:Bersacapavir PersScore 0.400 PoseScore 2.434
Pose 3:Bersacapavir PersScore 0.327 PoseScore 3.061
Pose 4:Bersacapavir PersScore 0.179 PoseScore 2.496
Pose 5:Bersacapavir PersScore 0.352 PoseScore 3.400
Pose 6:Bersacapavir PersScore 0.184 PoseScore 3.014
Pose 7:Bersacapavir PersScore 0.118 PoseScore 2.889
Pose 8:Bersacapavir PersScore 0.212 PoseScore 3.557
Pose 9:Bersacapavir PersScore 0.218 PoseScore 3.613
Pose 10:Bersacapavir PersScore 0.098 PoseScore 3.104
Pose 11:Bersacapavir PersScore 0.177 PoseScore 3.685
Pose 12:Bersacapavir PersScore 0.025 PoseScore 2.970
Pose 13:Bersacapavir PersScore 0.255 PoseScore 4.607
Pose 14:Bersacapavir PersScore 0.082 PoseScore 4.095
Pose 15:Bersacapavir PersScore 0.023 PoseScore 4.353
Pose 16:Bersacapavir PersScore 0.000 PoseScore 4.253
Pose 17:Bersacapavir PersScore 0.107 PoseScore 4.931
Pose 18:Bersacapavir PersScore 0.030 PoseScore 4.709
Pose 19:Bersacapavir PersScore 0.055 PoseScore 5.074
Pose 20:Bersacapavir PersScore 0.025 PoseScore 4.931
Pose 21:Bersacapavir PersScore 0.042 PoseScore 5.095
Pose 22:Bersacapavir PersScore 0.018 PoseScore 5.151
Pose 23:Bersacapavir PersScore 0.052 PoseScore 6.711
CV RMSD (A)
MD Time (ns)

**Figure S8. Binding pose metadynamics (BPMD) results for Arc1 and ARC complexes with Lenacapavir and Bersacapavir. Related to Figure 6. (a)** Arc1–Lenacapavir. **(b)** ARC–Lenacapavir. **(c)** Arc1–Bersacapavir. **(d)** ARC–Bersacapavir. Each panel summarizes the BPMD evaluation of representative poses obtained from IFD, with 20, 26, 28, and 23 poses analyzed for panels (a–d), respectively.
